# The biodiversity benefit of native forest over Grain-for-Green plantations

**DOI:** 10.1101/437418

**Authors:** Xiaoyang Wang, Fangyuan Hua, Lin Wang, David S. Wilcove, Douglas W Yu

## Abstract

**Aim:** China’s Grain-for-Green Program (GFGP) is the largest reforestation program in the world and has been operating since 1999. The GFGP has promoted the establishment of tree plantations over the preservation of diverse native forest. In a previous study (Hua et al. 2016, *Nat Comms* 7:12717), we showed that native forest supports higher species richnesses of birds and bees than do GFGP plantations. We also showed that ‘mixed-plantation’ GFGP plantations, which are mostly made up of two to five neighboring monoculture stands of different tree species, planted in checkboard fashion, support a level of bird (but not bee) species richness that is higher than any of the individual GFGP monocultures, although still below that of native forest. To better protect terrestrial biodiversity, which is an important objective of China’s land-sustainability spending, we recommended that the GFGP should firstly prioritize native forest conservation and regeneration and secondly promote checkerboard planting arrangements over monocultures. Here, we use metabarcoding of arthropod biodiversity to test the generality of these results and policy recommendations.

**Location:** Sichuan, China

**Methods:** We used COI-amplicon sequencing (‘metabarcoding’) of bulk samples of arthropods that were collected with pan traps in native forest, cropland, mixed plantations, and monocultures.

**Results:** Native forest supports the highest overall levels of arthropod species richness and diversity, followed by cropland and mixed plantations, followed by bamboo monoculture, followed by the other monocultures. Also, the arthropod community in mixed plantations shares more species with native forest than do any of the monocultures. Together, these results show a biodiversity value of mixed plantations for arthropods that is higher than that previously indicated by bees alone.

**Main conclusion:** These results strengthen our original policy recommendations of (1) promoting the conservation and expansion of native forest and (2) promoting mixed-plantation arrangements. The value of this added metabarcoding-based analysis is that these policy prescriptions are now also based on a dataset that includes over 500 species-resolution taxa, ranging across the Arthropoda.

## 1 INTRODUCTION

An important challenge for conservation science is to quantify the biodiversity impacts of major policy initiatives, especially in regions undergoing large shifts in land-use change. Nowhere is this more true than in China, which combines a high level of native biodiversity (Tao, Huang, Jin, & Guo, 2010) with a human population that is large and increasing its ecological footprint (Liu & Diamond, 2005; Pyne, 2013; Sayer & Sun, 2003; Xie et al., 2012). Moreover, China has had for decades the managerial, political, and financial capacity to implement the largest land-sustainability programs ever seen, from nature-reserve protection to reforestation to de-desertification (Bryan et al., 2018; Liu et al., 2003; Xu, Wang, & Xue, 1999). These programs have caused major land-use changes and successfully slowed land degradation caused by economic activities (Liu, Li, Ouyang, Tam & Chen, 2008; Ouyang et al., 2016; Ren et al., 2015). For example, China established its first nature reserve in 1956 and reached 2740 reserves at the end of 2015 (Ma, Shen, Grumbine, & Corlett, 2017). Nearly two-thirds of the area of those nature reserves have national-level status, meaning that they receive the highest level of protection and funding (Ren et al., 2015). National-level nature reserves have been shown to deter deforestation (Ren et al., 2015).

Two other major land-sustainability programs are the Natural Forest Protection Program (NFPP, also known as Natural Forest Conservation Program) and the Grain for Green Program (GFGP, also known as the Sloping Land Conservation Program and the Farm to Forest Program), which were implemented after widespread flooding in 1998 (Liu et al., 2008; Xu, Yin, Li, & Liu, 2006; Yin, Yin, & Li, 2009). The NFPP aims to reduce soil erosion and flooding by protecting native forests in the upstream watersheds of the Yangtze and Yellow Rivers (Liu et al., 2008; Ren et al., 2015). The GFGP complements the NFPP by controlling soil erosion on sloping land. The government pays cash and grain to farmers in exchange for tree planting on sloping farmland (Delang &Yuan, 2015; Liu et al., 2008; Ma et al., 2017; Xu et al., 2006; Zhai, Xu, Dai, Cannon, & Grumbine, 2014).

However, relative to their scale and budgets, little is known about the biodiversity consequences of China’s land-sustainability programs, even though an important goal of those programs has been promoted biodiversity conservation. In a recent, massive review, Bryan et al. (2018) were able to cite only one study on the consequences of reforestation programs for animal populations, Hua et al. (2016), although there is considerably more information on socioeconomic impacts (Liu et al., 2008; Liu & Lan, 2015; Long et al., 2006; Shen, Liao, & Yin, 2006; Yin, Liu, Zhao, Yao, & Liu, 2014) and basic ecosystem variables like water and soil maintenance, changes to vegetation coverage and structure, carbon storage, and so on (Deng, Liu, & Shangguan, 2014; Deng, Shangguan, & Li, 2012; Wang, Peng, Zhao, Liu, & Chen, 2017; Wang, Jiao, Rayburg, Wang, & Su, 2016; Wei et al., 2014; Zhou, Van Rompaey, & Wang, 2009). A series of reports have also focused on the replacement of native forest by plantations and the status of a few endangered species like giant panda (Hua et al., 2018; Liu et al., 2008; Ma, 2015; Ren et al., 2015; Wei et al., 2015; Zhai et al., 2014; Zheng & Cao, 2014).

Hua et al. (2016) surveyed bird and bee communities in GFGP forests in western Sichuan, comparing native forest remnants to GFGP-financed forest types, which include monoculture stands of bamboo, eucalyptus, and Japanese cedar, as well as ‘mixed plantations,’ which are patchworks (checkerboards) of two to five different monocultures and, to a lesser extent, *bona fide* tree-level mixtures (Hua et al., 2018). Most importantly, this study documented that bird and bee diversities were higher in native forest than in any of the monocultures. Or in other words, the biodiversity value of GFGP monocultures is low.

A secondary, and slightly surprising, result was that in mixed plantations, (non-breeding) bird species diversity was higher than in any of the individual monocultures, albeit lower than in native forest. In contrast, bee diversity was equally low in mixed plantations and monocultures. The lack of a biodiversity boost to bee diversity in mixed plantations was not surprising, since the understory vegetation in the monocultures was notably lacking in flowering plants (Hua et al., 2016). But this raises the question of why bird diversity was increased just by planting monocultures of different tree species next to each other. One possibility that could not be investigated in Hua et al. (2016) is that general arthropod diversity could also be boosted in the mixed plantations, since, unlike bees, other arthropods can exploit a range of food resources available even in plantations, via direct consumption of plants and fungi, and via decomposition, parasitism, and predation of other animals, including other arthropods. In turn, increased arthropod diversity might support more bird species.

The purpose of this study is to test the generality of the above results by interrogating the ‘rest of the biodiversity’ that was captured in the same sites analyzed by Hua et al. (2016). We employ the technique of metabarcoding, which combines traditional DNA barcoding with high-throughput DNA sequencing to characterize the biodiversity of mixed samples of eukaryotes (Cristescu, 2014; Deiner et al., 2017; Yu et al., 2012). A taxonomically informative genetic marker (here, a fragment of the mitochondrial cytochrome oxidase subunit I gene, which is used as the animal ‘DNA barcode’, Ratnasingham & Hebert, 2007) is polymerase-chain-amplified (PCR’d) from bulk samples, sequenced in parallel (here, on an Illumina sequencer), and processed bioinformatically to recover a sample X species table, where species are approximated by clusters of similar DNA sequences known as Operational Taxonomic Units (OTUs). These OTUs are matched against online databases (here, the Midori database (Machida, Leray, Ho, & Knowlton, 2017)) and assigned taxonomies. Finally, the tables are analyzed statistically to estimate community-ecology metrics, such as species diversity and turnover. Many bioinformatic pipelines are now available to carry out metabarcoding, and we describe ours below, also providing our scripts and data for replication. Metabarcoding has been shown to be a reliable and efficient method for biodiversity characterization (Ji et al., 2013).

We metabarcoded the non-bee arthropods caught by the same pan traps that we had used to trap bees, and we find that (1) native forest does support higher levels of arthropod species richness and diversity than do any of the GFGP-funded plantations (mixed plantations and monocultures) and also that (2) mixed plantations support significantly or marginally significantly higher levels of arthropod species richness and diversity than do the three individual monocultures: bamboo, Japanese cedar, and eucalyptus, paralleling the previous results with the birds, but not the bees. We also estimated phylogenetic diversity to test the robustness of our DNA-barcode clustering algorithm and found roughly the same results. We then visualized community compositions of all the forest types and found that mixed-plantation stands are (1) not surprisingly, made up of subsets from the three monocultures, and (2) somewhat surprisingly, compositionally most similar to native forest: mixed plantations uniquely share more species with native forest than does any of the individual monocultures. Beta diversity is dominated by species turnover, which means that even the monocultures support small but distinct arthropod communities, exemplifying Lindenmayer et al.’s (2006) insight that “management for diversity calls for diversity of management.” Overall, our results support the conclusions of Hua et al. (2016) that native forests contain the most biodiversity and should be prioritized in the GFGP for protection and expansion and that promotion of mixed planting of monocultures is a worthwhile but second-best policy reform for the GFGP.

## 2 METHODS

### 2.1 Study location

The study region and sampling locations are the same as those in Hua et al. (2016), where further details can be found. In short, our study region was a 7,949 km2 area in south-central Sichuan province (Figure 1) spanning an elevation range of 315-1,715 m above sea level, historically forested and then heavily deforested starting in the 1950s. The GFGP established ~54,800 ha of forest between 1999 and 2014, dominated by the following four types: short-rotation (6-20 years) monocultures of bamboo (BB), eucalyptus (EC), and Japanese cedar (JC), and compositionally simple, short-rotation (4) mixed plantations (MP) consisting of two to five tree species (including the three monoculture plantations). Monocultures are created by households planting the same tree species in neighboring stands of median 0.4 ha size. Correspondingly, mixed plantations are, in most cases, created by households planting different species in neighboring stands, resulting in a checkerboard structure, although about a quarter of mixed plantations do consist of individual-level mixtures. In Hua et al. (2016), we used the term ‘mixed forests’, but in Hua et al. (2018), we switched to ‘mixed plantations’ to emphasize that this forest type is man-made.

**Figure 1.**
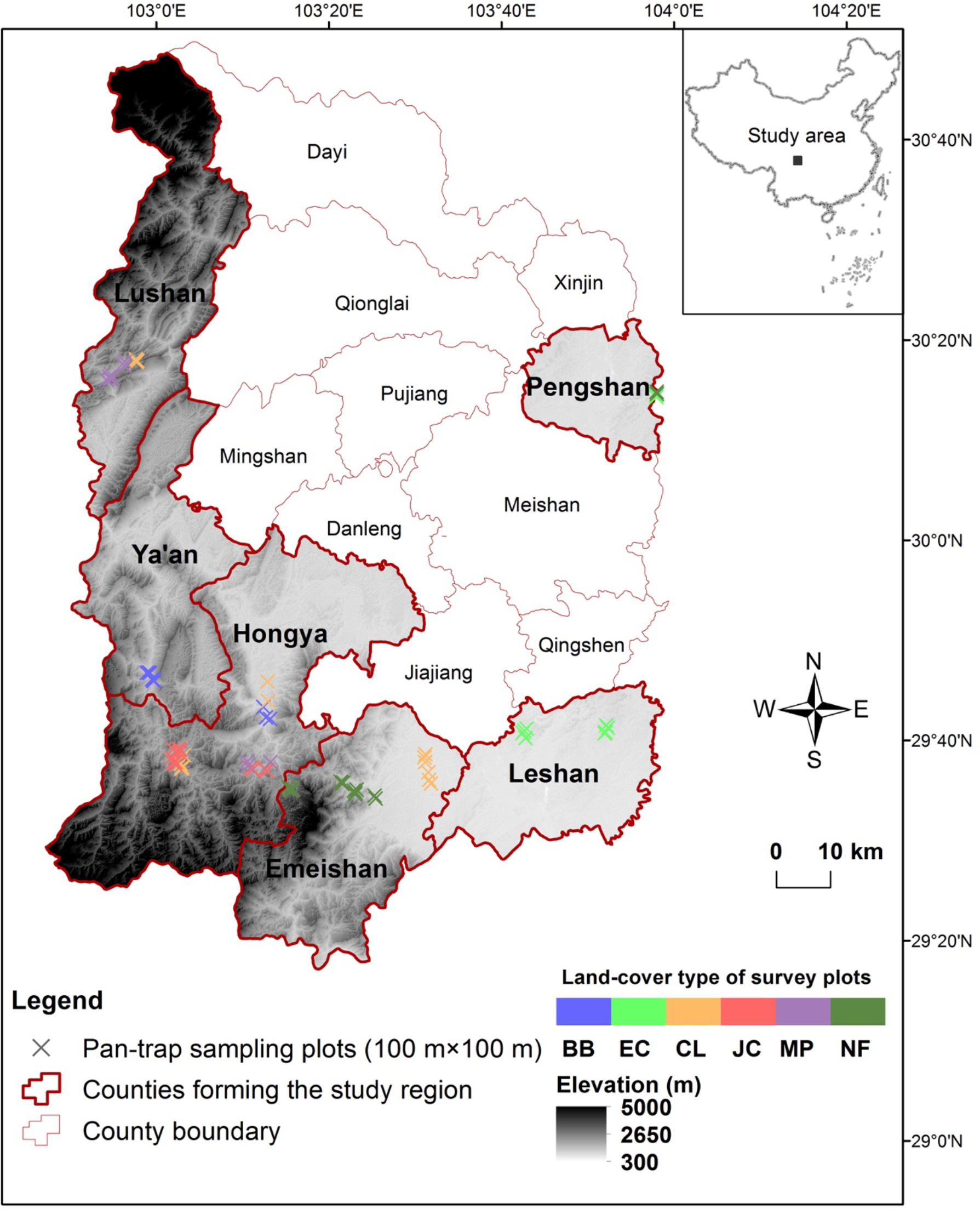
Study area in south-central Sichuan province, subdivided into counties and shaded by elevation. Each cross represents a pan-trap sampling location, color-coded by land-cover type: BB = bamboo monoculture, blue; EC = eucalyptus monoculture, light green; CL = cropland, orange; JC = Japanese cedar monoculture, red; MP = mixed plantations, purple; NF = native forest, dark green.

The two other surveyed habitats were cropland (CL) and native forest (NF). Cropland mostly consists of low-intensity plantings of rice, corn, and vegetables and is the land type that has been most displaced by GFGP forest. Native forest is broadleaf, subtropical, evergreen forest that has been subject to selective logging and other forms of extraction, especially since the 1950s. In part because this region of China has been inhabited for millennia, we were not able to locate any undisturbed native forest. Cropland is typically located on flatter land than are the forest plots, since GFGP reforestation targeted sloped land, and the native forests are concentrated toward the southern end of the study region, near Emei Mountain. For sampling, we chose larger expanses (> 60 ha) of each of these six land-cover types: BB, EC, JC, MP, NF, CL.

### 2.2 Sampling design

Each land-cover type was represented by at least two locations set ≥15 km apart. All forest stands chosen were closed canopy. For each land-cover type, we sampled with at least ten one-ha quadrats, within each of which we operated 40 fluorescent pan traps for 24 hrs (Bartholomew & Prowell, 2005) (Figure S1). In total, 74 quadrats were sampled (BB: n = 10 quadrats, EC: 10, JC: 12, MP: 10, NF: 16, CL: 16). Different quadrats were separated by ≥300 m if placed in the same forest stand. All individual samples were stored in 100% ethanol and stored at ambient temperature until shipment back to the Kunming Institute of Zoology, where they were stored in a −20 °C freezer before DNA extraction. The original reason for using pan traps had been to trap bees, which we DNA-barcoded individually for analysis in Hua et al. (2016). Here we use metabarcoding to analyze the pan-trap bycatch.

### 2.2 Amplicon preparation

For each of the 74 quadrats, we first pooled all 40 pan traps into a single sample. In this part, three quadrats had very few individuals, and we pooled them with their nearest-neighbor quadrat of the same land-cover type (EC02 and EC03 were pooled with EC01, NF03 was pooled with NF02). Thus, we were left with 71 bulk samples. Storage ethanol was removed by air drying on single-use filter papers. Our samples were dominated by Diptera and Hymenoptera, as expected with pan traps. We equalized input template DNA across species by using one leg of every individual larger than a mosquito and the whole body if smaller (e.g. midges). This was to reduce the effect of large-biomass individuals outcompeting small-biomass individuals during PCR, which improves taxon detection (Elbrecht, Peinert, & Leese, 2017). DNA extraction followed the protocols of Qiagen DNeasy Blood & Tissue Kits (Hilden, Germany), followed by quantification via Nanodrop 2000 spectrophotometer (Thermo Fisher Scientific, Wilmington, DE).

We amplified a 319-bp fragment of COI using forward primer LCO1490 (5’-GGTCAACAAATCATAAAGATATTGG-3’) and reverse primer mlCOIintR (5’-GGNGGRTANANNGTYCANCCNGYNCC-3’) (Leray et al., 2013). All samples were carried out with two rounds of PCR. In the first round, both forward and reverse primers (12-17 bp) were tailed with tags to allow sample identification. In the second-round PCR, we added Illumina adapters to the amplicons produced in the first PCR, thus avoiding the possibility of tag jumping that can arise by carrying out library preparation on mixtures of amplicons (Schnell, Bohmann, & Gilbert, 2015). A table of the primers with tags and the second-round PCR primers is in Supplementary Information (Table S1). All PCRs were performed on a Mastercycler Pro (Eppendorf, Germany) in 20-μl reaction volumes, each containing 2 μl 10 × buffer (Mg^2+^ plus), 0.2 mM dNTPs, 0.4 μM of each primer, 1 μl DMSO, 0.4 μl BSA (bovine serum albumin) (TaKaRa Biotechnology Co. Ltd, Dalian, China), 0.6 U exTaq DNA polymerase (TaKaRa Biotechnology), and approximately 60 ng genomic DNA. Both rounds of PCR started with an initial denaturation process at 94 °C for 4 mins, followed by 35 cycles of 94 °C for 45s, 45 °C for 45s, 72 °C for 90s, and finishing at 72 °C for 10 mins. PCR products were gel-purified with QIAquick PCR Purification Kit (Qiagen). One sample failed to amplify. For the remaining 70 samples, we pooled purified PCR products into two libraries, then sequenced on the Illumina MiSeq platform (Reagent Kit v3 600 cycles, 300PE) at the Southwest Biodiversity Institute Regional Instrument Center in Kunming. The total number of paired-end sequences returned was 13,601,908.

### 2.3 Data analyses

The bioinformatic script (including parameter values) for the analyses below are provided in Supplementary Information and will also be archived in datadryad.org, along with the raw sequence data and metadata. The *R* scripts and data tables are on https://github.com/dougwyu/Sichuan2014.

#### 2.3.1 Bioinformatic processing

##### Initial processing

We first removed remnant Illumina adapter sequences from the MiSeq output files with *AdapterRemoval* 2.2.0 (Schubert, Lindgreen, & Orlando, 2016), followed by Schirmer et al.’s (2015) recommended pipeline to filter, trim, denoise, and merge read pairs. Specifically, we trimmed low-quality ends using *sickle* 1.33 (Joshi & Fass, 2011), corrected sequence errors using the *BayesHammer* module in *SPAdes* 3.10.1 (Nikolenko, Korobeynikov, & Alekseyev, 2013), and merged reads using *PandaSeq* 2.11 (Masella, Bartram, Truszkowski, Brown, & Neufeld, 2012). In all cases, we used default parameters.

##### Demultiplexing and Clustering

We then used *qiime* 1.9.1’s *split_libraries.py* function (Caporaso et al., 2010) to demultiplex reads by samples, and we used *usearch* 9.2.64 (Edgar, 2010) to keep only reads between 300 and 330 bp, inclusive, since our target amplicon is 319 bp. We used *vsearch* 2.4.3 (Rognes, Flouri, Nichols, Quince, & Mahé, 2016) to carry out *de-novo* chimera removal and clustered the reads into 3,507 97% Operational Taxonomic Units (OTUs) using *CROP* 1.33 (Hao, Jiang, & Chen, 2011). We also tried *swarm* 2.2.2 (Mahé, Rognes, Quince, de Vargas, & Dunthorn, 2015), but it returned huge numbers of OTUs that could not be reduced even after running through ‘lulu’ (see below).

##### OTU filtration and taxonomic assignment

From the resulting OTU table (i.e. sample X ‘species’ table), we used the *R* package ‘lulu’ 0.1.0 (Frøslev et al., 2017) to combine OTUs that are likely from the same species but which CROP failed to cluster. ‘lulu’ infers (and combines) such ‘parent-child’ sets by first calculating pairwise similarities of all OTU representative sequences (here, using *vsearch*) to identify sets of high-similarity OTUs and then combining OTUs within such sets that show nested distributions across samples. For example, four OTUs might show high pairwise similarities, and within this set of four, one OTU contains the most reads and is observed in ten samples. This OTU is designated the parent, and daughter OTUs are inferred if they are only present in a subset of the parent OTU’s ten samples. We obtained 1,506 OTUs at the end of this step.

A common filtering step is to remove OTUs made up of few reads (e.g. 1-read OTUs), as these are more likely to be artefactual. For instance, PCR errors can generate clusters of sequences that form their own OTUs and are sufficiently different from the parent that they cannot be identified as daughters by ‘lulu’. Such OTUs are more likely to be small because these novel haplotypes typically arise in a later PCR cycle and are thus amplified less often than the original true haplotypes. There is no reason for such PCR errors to be sequenced at low quality, so they cannot be filtered out by quality score. However, the definition of ‘few reads’ is inherently subjective and necessarily differs with the size of the sequence dataset (and other aspects of the lab and bioinformatic pipeline). We therefore used ‘phyloseq’ 1.19.1 (McMurdie & Holmes, 2013) to plot the number of OTUs that would be filtered out at different minimum OTU sizes (see http://evomics.org/wp-content/uploads/2016/01/phyloseq-Lab-01-Answers.html, accessed 19 July 2018), and we chose a minimum OTU size of 44 reads, which is roughly the inflection point of the above graph and is the size that filters out the largest number of OTUs for the smallest OTU size minimum. This removed about 60% of the original 1,506 OTUs, and we ended this step with 594 OTUs.

We then used *PyNAST* 1.2.2 to align the 594 remaining OTU representative sequences to a reference alignment of Arthropoda COI sequences (from Yu et al. 2012) at a minimum similarity of 60%; one sequence failed to align and was deleted. The remaining sequences were translated to amino acids using the invertebrate mitochondrial codon table, and we removed 32 OTUs with representative sequences that contain stop codons. We carried out taxonomic assignment of the OTUs using the Naïve Bayesian Classifier (Wang, Garrity, Tiedje, & Cole, 2007) that had been trained on the Midori UNIQUE COI dataset (Machida et al., 2017). Sixteen OTUs assigned to non-Arthropoda taxa and two OTUs assigned to Collembola were removed. We ended this step with 543 OTUs.

Finally, we inspected the OTU table and set to zero those cells that had <5 reads representing that OTU in that sample, since these are more likely to be the result of sequencing error. In addition, we removed two rows (samples) that contained ≤100 reads total (i.e. two samples with little data due to sequencing or PCR failure) and removed seven rows (samples) with <5 OTUs because these samples are potentially overly influential (‘ecological outliers’) in analyses of species richness. These seven samples included 2 from native forest and 5 from monocultures (3 BB, 1 EC, 1 JC), meaning that we disproportionately removed very low-diversity samples from the monocultures, making our species diversity analyses below more conservative. After these sample removals, seven OTUs were left with <20 reads and were removed. Because we do not consider read numbers per OTU to be reliable measures of biomass or abundance (Nichols et al., 2018; Piñol, Mir, Gomez-Polo, & Agustí, 2015; Yu et al., 2012), we converted the OTU table to presence/absence (0/1). Throughout, our bias was to remove false-positive detections even at the expense of losing true-positive detections, thereby resulting in a dataset with less, but more reliable, data. We ended with 536 OTUs and 61 samples.

#### 2.3.2 Community analysis

##### Alpha diversity

All community analyses were performed in *R* 3.3.3 (R Core Team, 2017). We plotted observed species richnesses using ‘beanplot’ 1.2 (Kampstra, 2008) and estimated species richnesses and diversities using two sample-based estimators: function *specpool* in ‘vegan’ 2.4-5 (Chiu, Wang, Walther, & Chao, 2014) and ‘iNEXT’ 2.0.12 (Hsieh, H.Ma, & Chao, 2016).

Because we used a combination of *CROP*+‘lulu’ and ‘phyloseq’ to combine and remove, respectively, small OTUs that are likely to be artefactual, the OTUs that remain are more likely to represent true presences. Nonetheless, it remains possible that we have still over-split some biological species into multiple OTUs, since there is no single correct similarity threshold for species delimitation, and this oversplitting might have occurred more often for some taxa in some land-cover types, leading to artefactual differences in species richness. However, such oversplit OTUs should cluster together in a phylogenetic tree and thus contribute less to estimates of phylogenetic diversity than would OTUs from multiple, true biological species. Phylogenetic diversity should thus be a more robust estimator of alpha diversity (Yu et al., 2012). To estimate sample phylogenetic diversities, we used ‘iNextPD’ 0.3.2 (Hsieh & Chao, 2017). We built a maximum-likelihood (ML) tree in *RaxML* 8.0.0 (Stamatakis, 2014) with an alignment of the OTU representative sequences (using a General Time Reversible (GTR) model of nucleotide substitution and a gamma model of rate heterogeneity estimating the proportion of invariable sites (-m GTRGAMMAI)). The algorithm used a rapid bootstrap analysis and searched for the best-scoring ML tree (-f a), with 1000 times bootstrap (-N 1000) and 12345 set to be the parsimony random seed (-p 12345)). Three of the sequences produced very long branches in the ML tree, which would skew estimates of phylogenetic diversity, and we removed these three OTUS and their representative sequences from the ‘iNextPD’ analysis.

##### Beta diversity

To visualize community composition, we ran a Bayesian ordination using ‘boral’ 1.6.1 (Hui, 2016), which is more statistically robust than running non-metric multidimensional analysis (NMDS) because ‘boral’ is model-based and thus allows us to apply a suitable error distribution so that fitted-model residuals are properly distributed. We used a binomial error distribution and no row effect to fit the model since we were using presence/absence data (Figure S2). For the same reasons, we used ‘mvabund’ 3.12.3 (Wang, Naumann, Wright, & Warton, 2012) to test the hypotheses that the compositions of native forest and mixed plantations differ from each other and differ from the monocultures and cropland.

We also visualized beta diversity with an ‘UpSetR’ 1.3.3 intersection diagram (an easier-to-parse alternative to Venn diagrams (Conway, Lex, & Gehlenborg, 2017)) and with a heatmap using the *tabasco* function in ‘vegan’. We then partitioned the beta diversity into turnover and nestedness components using ‘betapart’ 1.4-1 (Baselga & Orme, 2012) with binary Jaccard dissimilarities, and we visually compared these two components using the *metaMDS* function in ‘vegan’. Finally, we used ‘metacoder’ 0.2.0 (Foster, Sharpton, & Grunwald, 2017) to generate taxonomic ‘heat trees’ (instead of barcharts) to visualize and pairwise-compare the taxonomic compositions of the six land-cover types, in order to evaluate whether beta diversity is being driven by specific taxa. The ‘metacoder’ analysis used the presence/absence OTU table.

## 3 RESULTS

### 3.1 Alpha diversity

#### Species diversity

After correction for multiple tests, *observed* species richnesses did not differ significantly between any pair of forest types (Figure 2), but observed species richnesses ignore undetected species. We thus first used the Chao2 estimator (Chao, 1987) and found that native forest, mixed plantations, and cropland have the highest estimated species richnesses and do not differ significantly in richness from each other (Figure 3).

**Figure 2.**
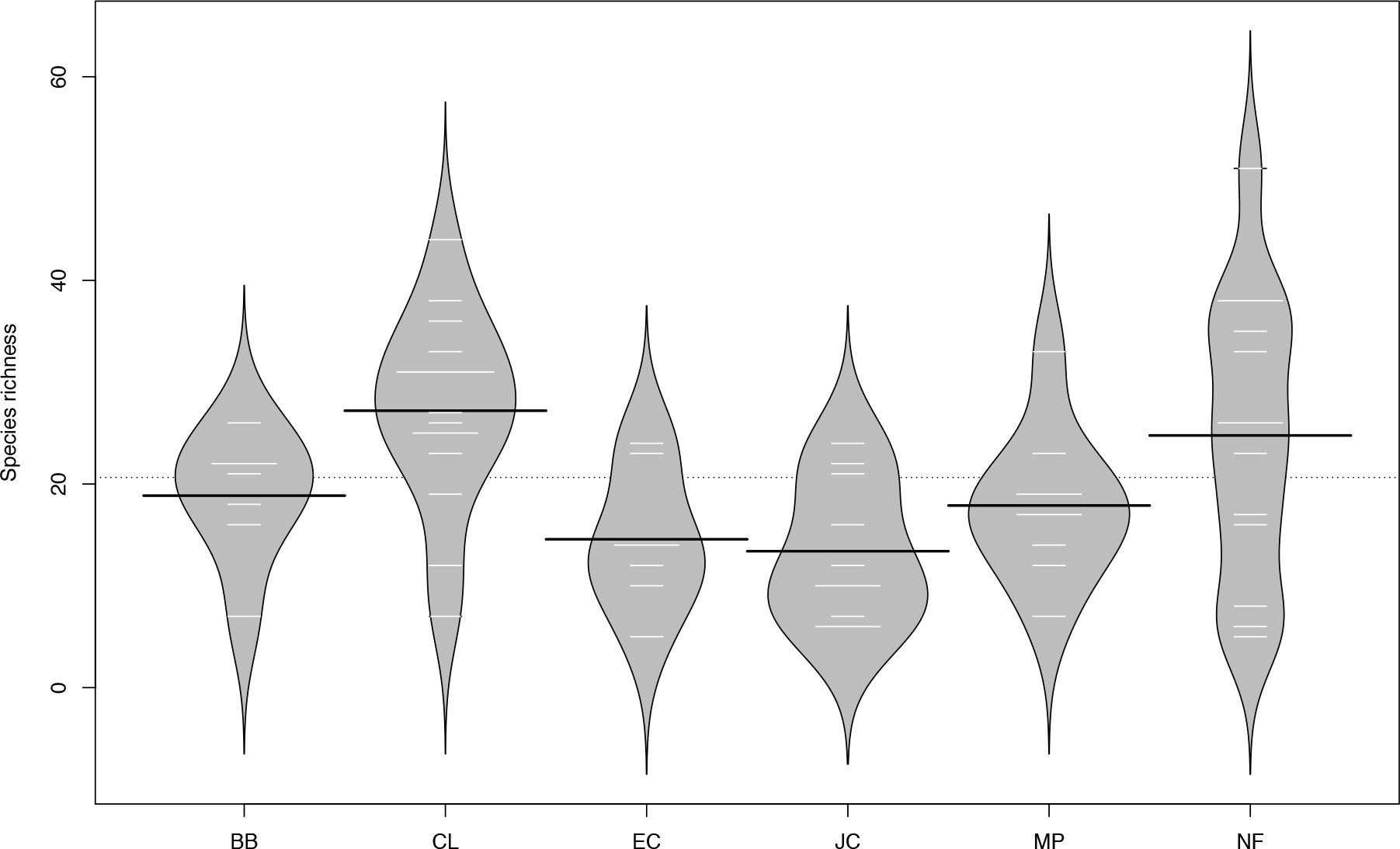
Beanplots of each habitat’s observed species richness. White lines are observed values at each sampling site, black lines are the mean per land-cover type, and the dashed line is the grand mean. Codes for land-cover types as in Figure 1.

**Figure 3.**
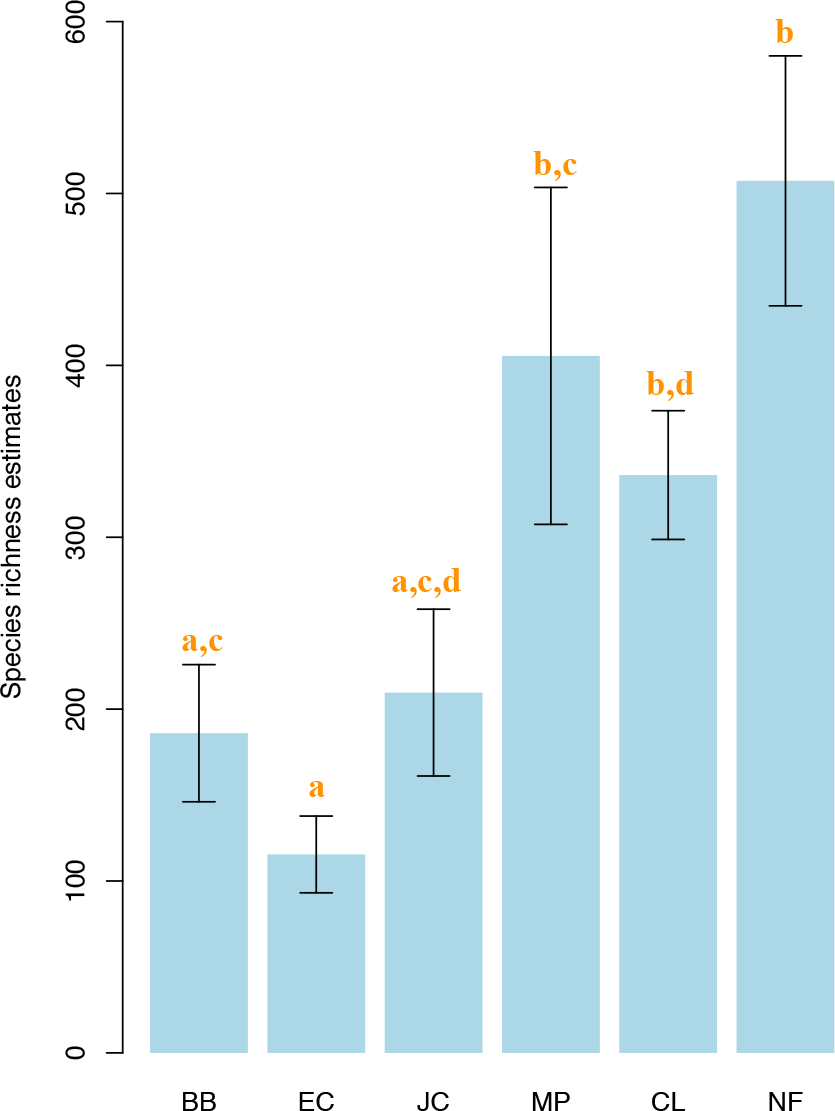
Comparisons of Chao2 species richness estimates across habitat type. Forest types sharing the same superscript are not significantly different at the p=0.05 level after table-wide correction for multiple tests (Welch’s t-test).

Importantly, the monocultures (bamboo, eucalyptus, and Japanese cedar) are estimated to support half or less of the species richnesses of native forests and mixed plantations, and after correction for multiple pairwise tests (`p.adjust(method=“fdr”)`), these differences achieve formal statistical significance for all comparisons with native forest and for the comparison between mixed plantations and eucalyptus forest (MP:EC, p = 0.046). The pairwise comparisons between mixed plantations and the other two monocultures achieve marginal significance (MP:BB, p = 0.10; MP:JC, p = 0.14).

We then checked the robustness of this finding with ‘iNEXT’ and ‘iNextPD’. As with the Chao2 estimator, ‘iNEXT’ found that native forest has the highest estimated asymptotic species richness and species diversities (Shannon and Simpson indices), followed by cropland and mixed plantations, followed by the three monocultures (Figure 4). Native-forest species richness and diversity are significantly higher than in all the other land-cover types (the confidence intervals do not overlap), and the richness and diversities of mixed plantations are significantly higher than all the monocultures, with the possible exception of bamboo (the MP and BB confidence intervals touch for species richness, do not touch for Shannon and Simpson diversities). We note that 95% confidence-interval overlap is considered an overly conservative test for significance at the p=0.05 level (MacGregor-Fors & Payton, 2013).

Using’ iNextPD’, the estimators *Faith’s phylogenetic diversity* and *Phylogenetic Entropy* show roughly the same ranking pattern (native forest, followed by cropland and mixed plantations, followed by the three monocultures, but with more confidence-interval overlap amongst mixed plantations, cropland, and bamboo) (Figure 5). The final estimator, Rao’s quadratic entropy, shows confidence-interval overlap of mixed plantations with two of the three monocultures, but the rank order of phylogenetic diversities is the same.

**Figure 4.**
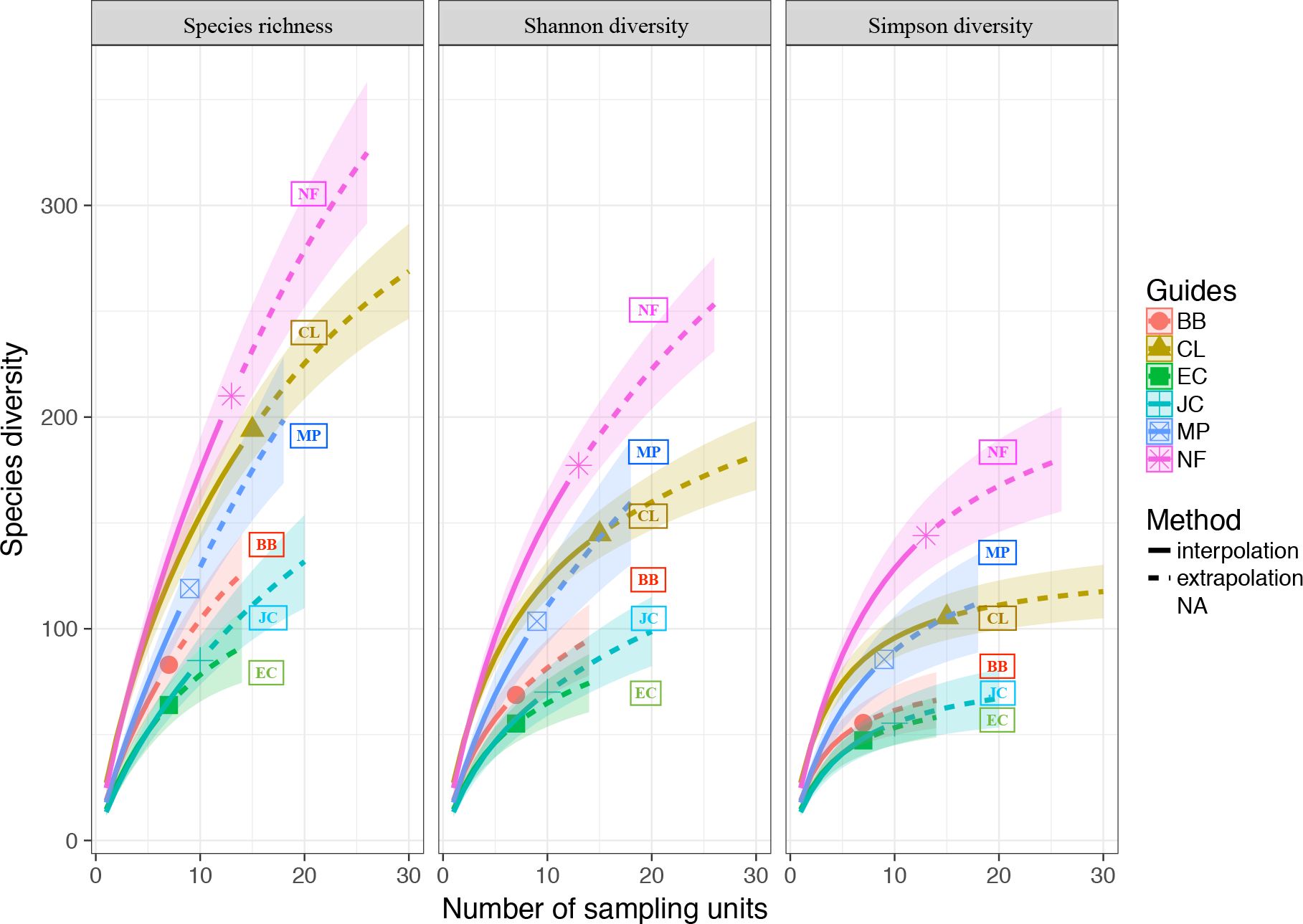
‘iNEXT’ estimates of species richness, Shannon diversity, and Simpson diversity by land-cover type, using sample-based rarefaction and extrapolation. Native forest (NF) has the highest species richness and diversities, followed by cropland (CL) and mixed plantation (MP), followed by the three monoculture plantations (BB, EC, and JC). Codes for land-cover types as in Figure 1. Symbols on each curve indicate the number of sampled locations per land-cover type, solid lines represent ‘iNEXT’ interpolations, and dashed lines represent ‘iNEXT’ extrapolations, with 95% confidence intervals. Statistically significant pairwise differences are detected visually by non-overlapping confidence intervals, and are somewhat conservative (MacGregor-Fors & Payton 2013).

**Figure 5.**
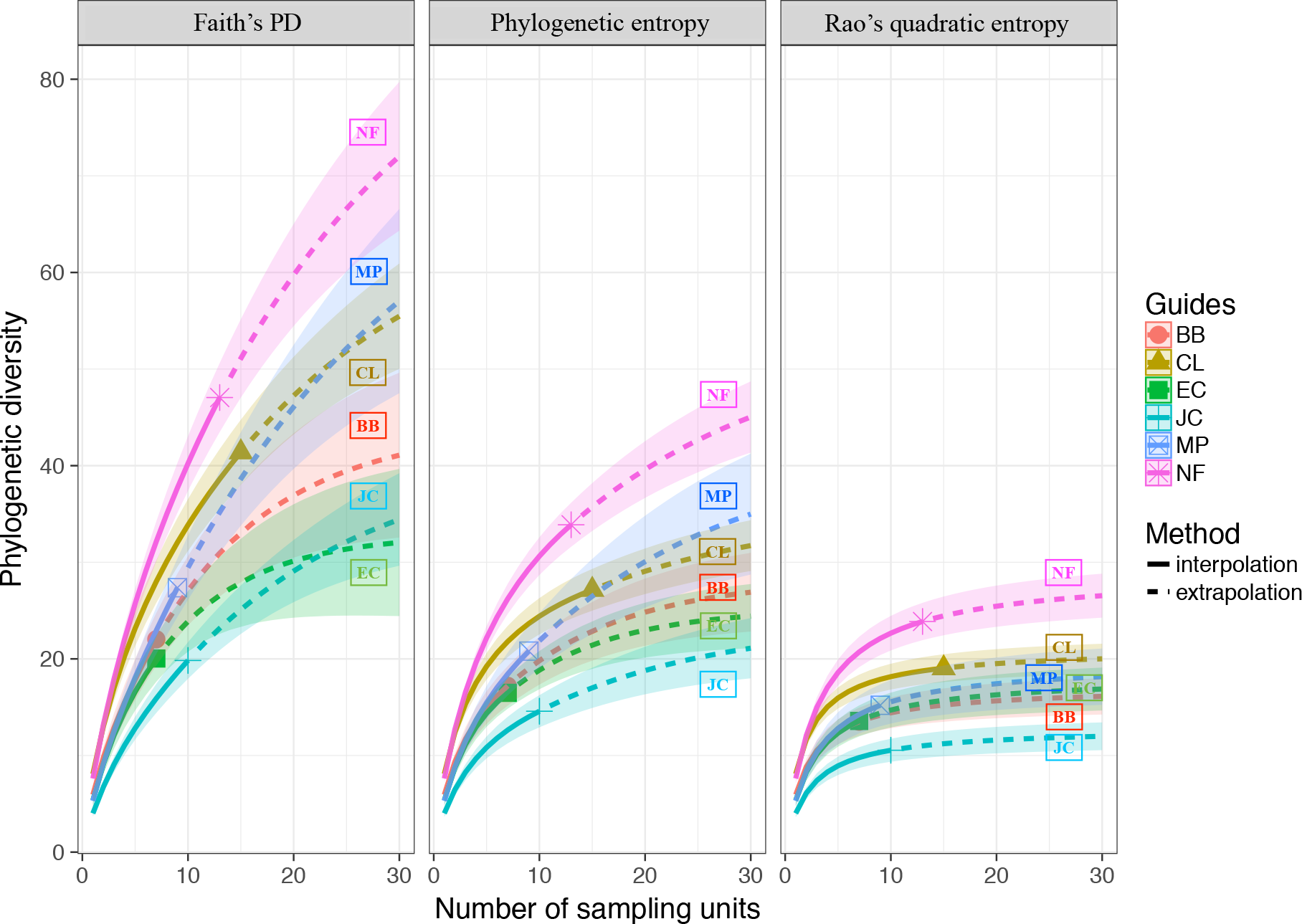
‘iNextPD’ estimates of phylogenetic diversity by land-cover type, using sample-based rarefaction and extrapolation. Similar to the results in Figure 4, two of the three estimators of phylogenetic diversity are higher in native forest (NF), followed by cropland (CL) and mixed plantation (MP), followed by the three monocultures (BB, EC, and JC; codes for land-cover types as in Figure 1). Symbols indicate sample sizes per land-cover type, solid lines represent ‘iNextPD’ interpolations, and dashed lines represent ‘iNextPD’ extrapolations, with 95% confidence intervals.

In summary, two incidence-based methods (Chao2, ‘iNEXT’) and one phylogenetic-diversity-based method (‘iNextPD’) all find that species richness and diversity in native forest is highest and that the mixed plantations are significantly or marginally significantly more diverse than the monoculture plantations (Japanese cedar, eucalyptus, bamboo).

We can also use ‘iNextPD’ to visualize phylogenetic coverage by land-cover type (Figure 6), and we see that the three monocultures, and the mixed plantations to a lesser extent, exhibit multiple coverage deficits, while cropland and native forest have almost complete coverage of the OTU tree. In the next section, we explore these compositional differences between habitats.

**Figure 6.**
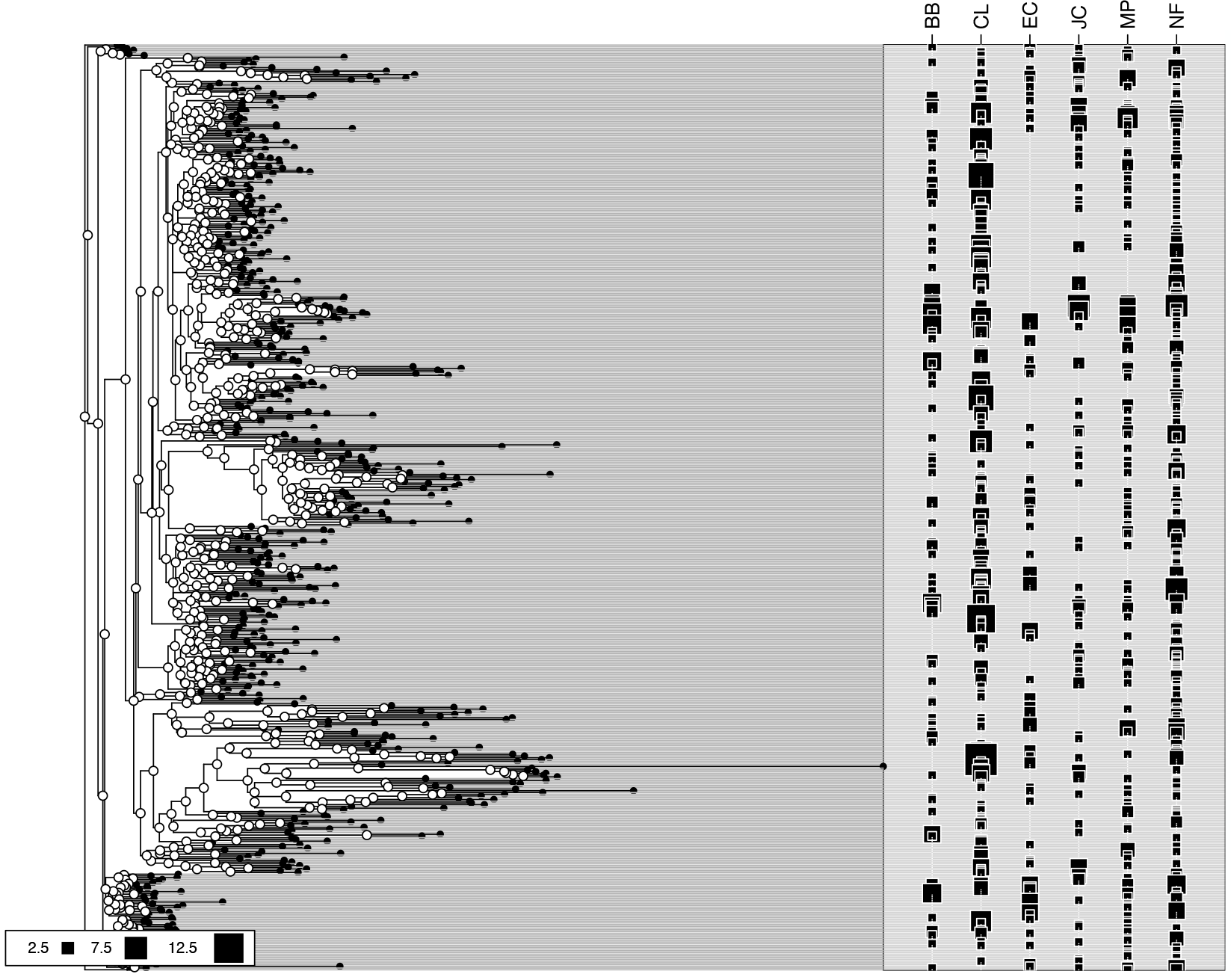
Phylogenetic distribution of OTUs by land-cover type, created using ‘iNextPD’. Terminal nodes are black and represent the OTUs. Internal nodes are white. Sizes of the squares on the right indicate each OTU’s incidence frequency (number of samples in which the OTU is observed). Phylogenetic coverage is most complete in native forest (NF) and cropland (CL), followed by mixed plantation (MP), followed by the three monocultures (BB, EC, JC). Codes for land-cover types as in Figure 1.

### 3.2 Beta diversity

We carried out a model-based, unconstrained ordination with ‘boral’ to visualize compositional differences amongst the six land-cover types (Figure 7). Not surprisingly, the primary separation was between cropland and the other land-cover types, which were arranged at different ends of the first latent-variable axis. The cropland sites themselves also clustered into two groups by elevation. A non-metric multidimensional scaling (NMDS) ordination showed a similar pattern but failed to separate cropland sites into high- and low-elevation clusters (data not shown).

**Figure 7.**
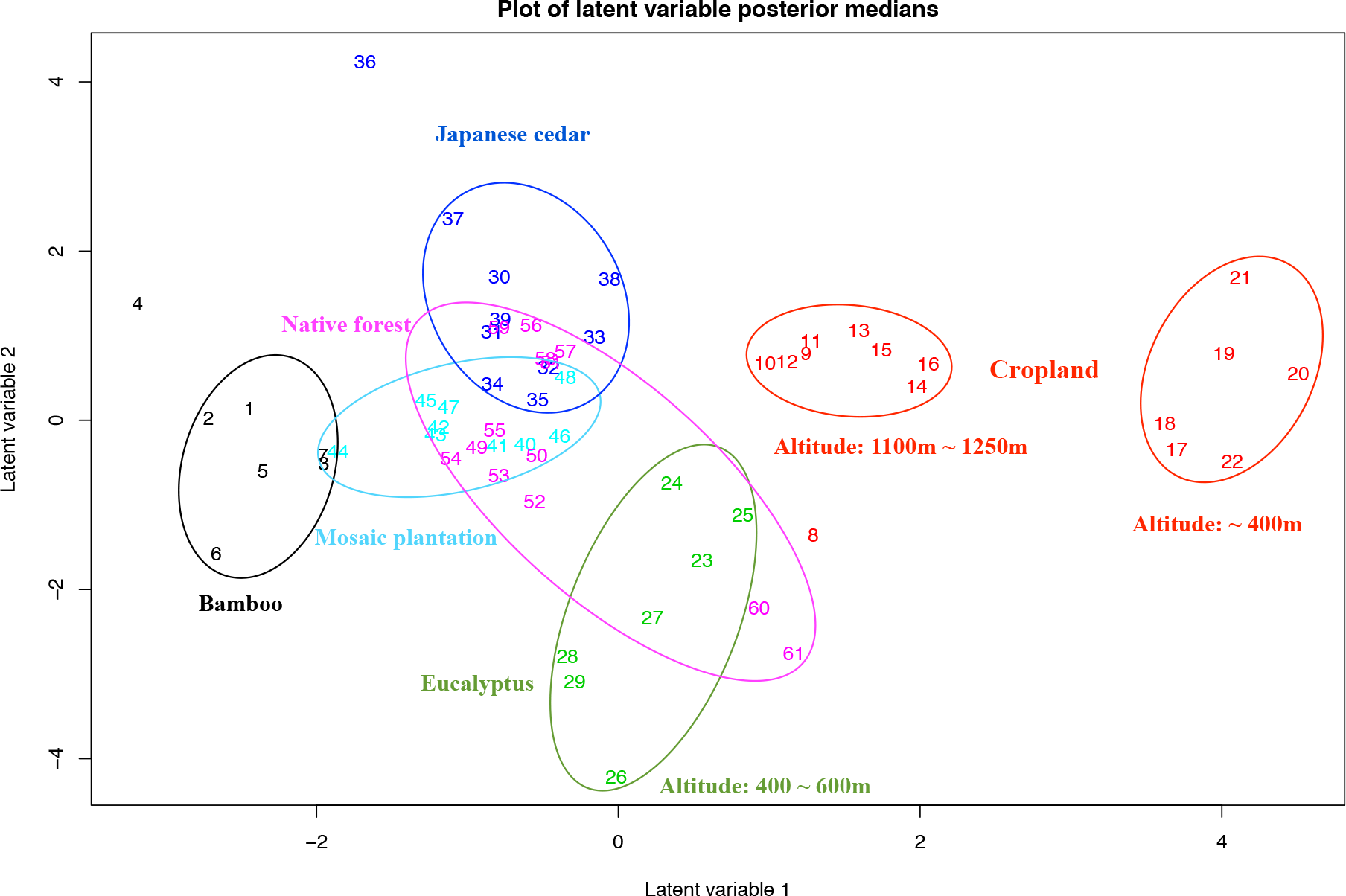
‘Boral’ ordination of beta diversity by land-cover type. Colors represent land-cover types, and numbers represent individual samples. Cropland (CL) sites separate into two clusters by elevation. Overlap of native forest (NF) and mixed-plantation (MP) points indicates greater compositional similarity between these two land-cover types. Ovals manually added to visualize community groupings. Residuals of the ‘boral’ fit in Fig. S2.

The latent variables that are output by ‘boral’ (Figure 7) can be thought of as ‘unmeasured predictor variables’ that are revealed by correlations amongst species in their distributions across samples (Warton et al., 2015). Latent variable 1 is correlated with elevation (r = −0.457, df = 59, p = 0.0002). Latent variable 2 largely separates eucalyptus monocultures from the other land-cover types, which might reflect eucalyptus’ distinct phytochemistry. Importantly, the mixed-plantation and (most of) the native-forest sites overlap each other and are encircled by the monocultures, indicating that native forest and mixed plantations are compositionally most similar.

The ‘UpSetR’ intersection diagram (Figure 8) provides a granular view of the above results. Firstly, consistent with what we found in the diversity analyses (Figures 3, 4, 5), native forest (110 OTUs) and cropland (130 OTUs) support more than 2.5 times the number of ‘unique species’ (species observed only in one land-cover type) than any of the plantations, and secondly, of the plantations, mixed plantations support the most unique species (44 OTUs). The greater compositional similarity that native forest has with mixed plantations (Figure 7) is partly explained by native forest uniquely sharing more OTUs with mixed plantations (22 OTUs) than with any of the monocultures (13, 9, and 5 OTUs).

**Figure 8.**
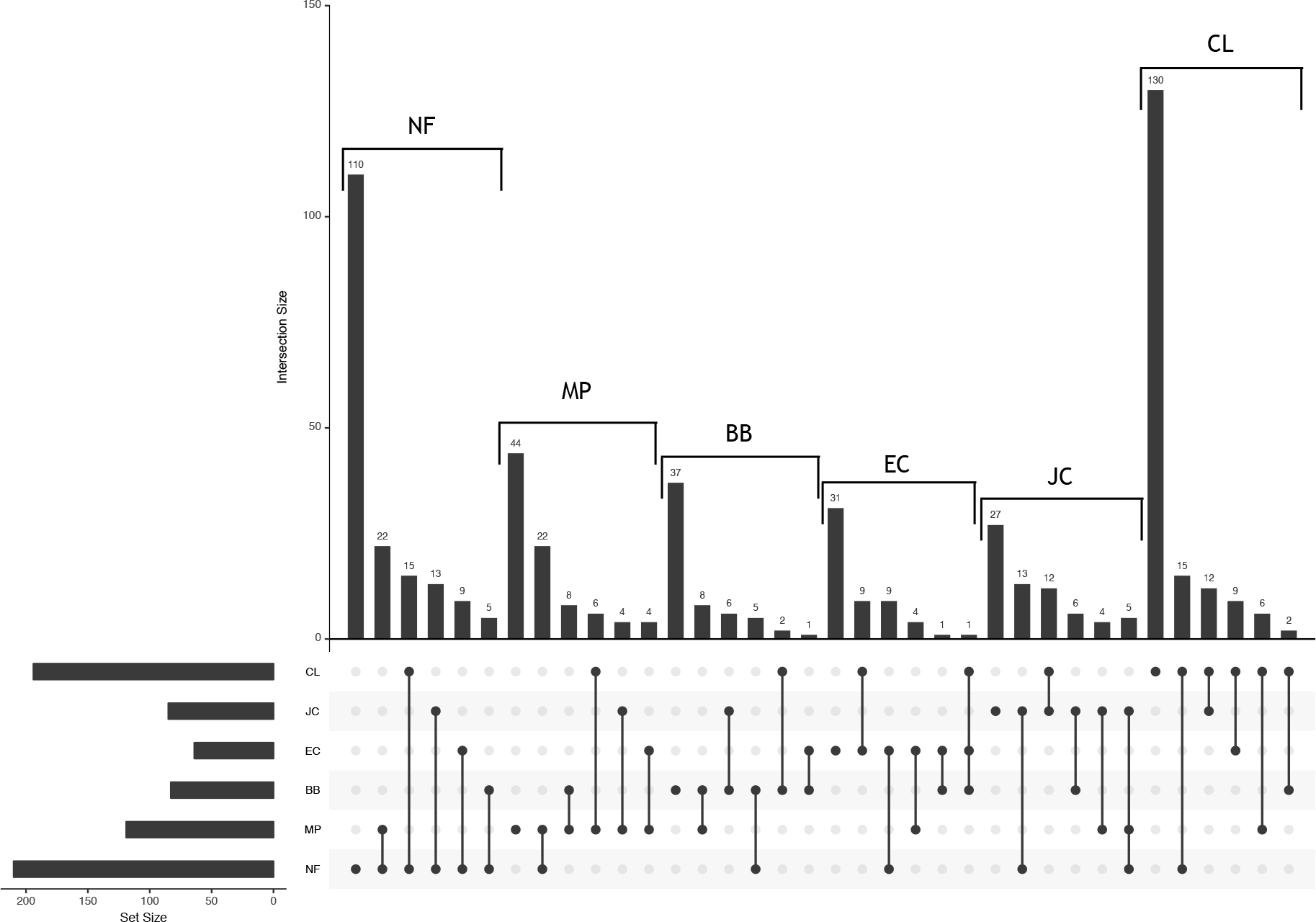
UpSetR intersection map of OTUs unique to and shared among land-cover types. Cropland (CL=130) and native forest (NF=110) support the most unique OTUs, followed by the four plantations (MP=44, BB=37, EC=31, JC=27). Native forest uniquely shares almost as many OTUs with Mixed plantations (22 OTUs) as native forest does with the three monocultures combined (27 OTUs, =13+9+5). Horizontal bars on the left indicate the number of OTUs in each land-cover class. Codes for land-cover types as in Figure 1. For clarity, only pairwise comparisons shown. A non-truncated version is presented in Fig. S3.

However, despite their overlap, *mvabund* analysis shows that the arthropod communities of mixed plantations and native forest are still significantly distinct from each other, and from the three monocultures and cropland (Table 1).

**Table 1.** *mvabund* compositional comparisons. We used *mvabund* to test whether arthropod species compositions in native forest (NF) and mixed plantations (MP) are significantly different from each other and from the other land-cover types in the study region: bamboo (BB), eucalyptus (EC), Japanese cedar (JC), and cropland (CL). After Bonferroni correction, all comparisons were significantly different at *p* < 0.01.

|  | MP | BB | CL | EC | JC |
| --- | --- | --- | --- | --- | --- |
| NF | 0.00125 | 0.00125 | 0.00125 | 0.00125 | 0.003 |
| MP |  | 0.001 | 0.001 | 0.001 | 0.001 |

#### Turnover versus nestedness

We now ask whether the compositional differences amongst land-cover types are driven by turnover or nestedness. ‘betapart’ analysis shows that turnover, not nestedness, dominates compositional differences (Figure 9), which is consistent with the UpSetR diagram showing that the mode OTU category in each land-cover type is unique OTUs. In other words, the monoculture arthropod communities are not just subsets of the richer native-forest, cropland, and mixed-plantation arthropod communities but rather contain their own monoculture-specific sets of species. We provide a heatmap in supplementary information (Figure S4) as another visualization of the dominance of turnover.

**Figure 9.**
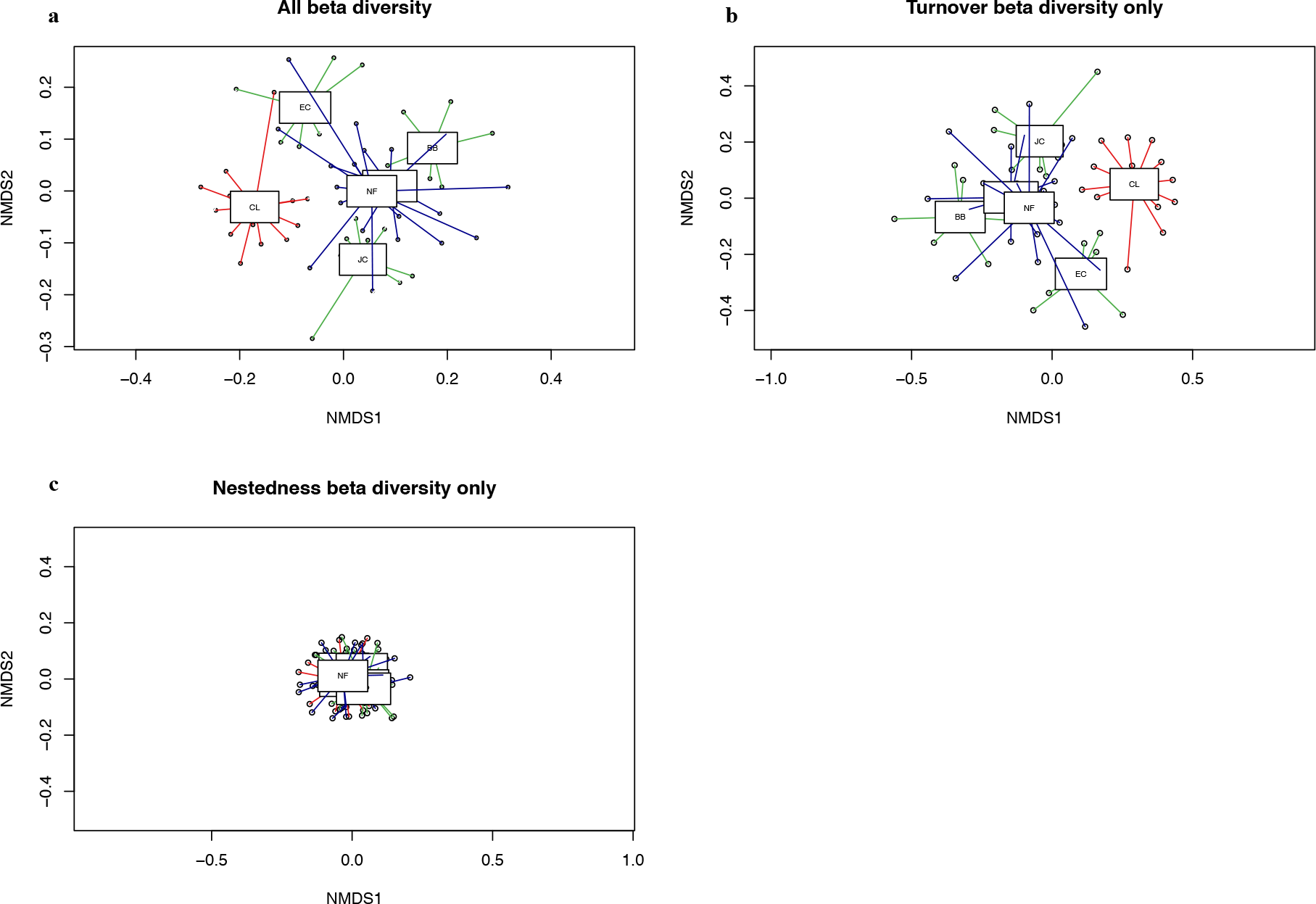
NMDS (non-metric multidimensional scaling) ordination of beta diversity by habitat type (binary Jaccard dissimilarities), partitioned with ‘betapart’. a. Total beta diversity. b. Beta diversity based on species turnover only. c. Beta diversity based on species nestedness only. Turnover accounts for most the observed beta diversity across habitats. Codes for land-cover types as in Figure 1.

#### Taxonomic compositions of and differences between land-cover types

The 536 arthropod species in our metabarcoding dataset represent a wide range of arachnid and insect orders and thus, represent a wide range of ecological functions (Figure 10), including generalist predators (Araneae, Formicidae) and more specialized parasites and parasitoids (Tachinidae, Phoridae, Braconidae) of other arthropods. Lower down the trophic level, we observe taxa that are noted for pollination (Thysanoptera, Syrphidae), xylophagy (Isoptera), and various modes of detritivory, fungivory, frugivory, herbivory, and animal parasitism (Lepidoptera, Hemiptera, Diptera, Orthoptera, Formicidae, Thysanoptera).

**Figure 10.**
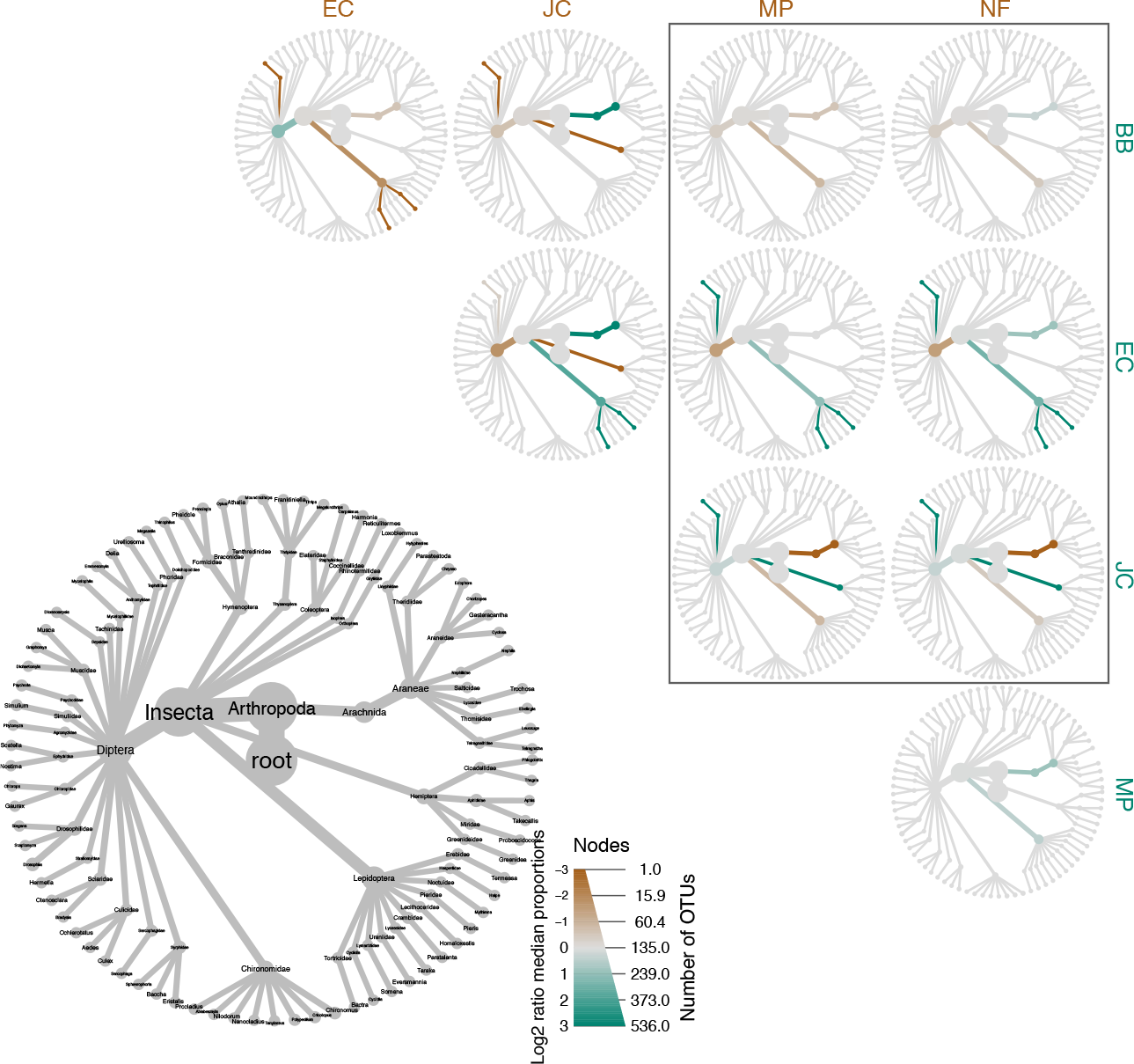
Pairwise taxonomic comparisons of all land-cover types. Upper right triangle: greener branches indicate taxa that are relatively more abundant (in terms of numbers of OTUs) in the land-cover type along the right column, and browner branches indicate taxa that are relatively more abundant in the land-cover type along the top row. Lower left: taxonomic identities of the branches. Note that this is a taxonomic tree, not a phylogenetic tree. Legend: width indicates number of OTUs at a given taxonomic rank, and color indicates relative differences in log_2_(number of OTUs). Codes for land-cover types as in Figure 1. A figure including cropland is in supplementary information (Figure S5).

The ‘boral’ ordination (Figure 7) reveals compositional similarity between mixed plantations and native forests, but which taxa are most responsible for this similarity, and for the differences with the other forest types? The ‘metacoder’ heat trees (Figure 10, inset box) show that mixed plantations and native forests *differ in the same ways* from each of the three monocultures. Relative to bamboo, mixed-plantation and native-forest sites both have slightly more Lepidoptera-assigned OTUs. Relative to eucalyptus, mixed-plantation and native-forest sites both have more Diptera-assigned OTUs and fewer of three OTUs assigned to genera *Mycetophila*, *Sonema*, and *Homaloxestis*, which can be taken as eucalyptus indicator species. Finally, relative to Japanese cedar, mixed-plantation and native-forest sites both have more Araneae- and Lepidoptera-assigned OTUs, fewer Hemiptera-assigned OTUs, and fewer of the OTU assigned to *Mycetophila*. Heat-tree differences at higher taxonomic ranks (e.g. more Araneae-assigned OTUs) mean that the OTUs that separate two land-cover types differ across sample pairs but nonetheless are in the Araneae and Diptera.

Finally, and not surprisingly, when we include cropland in the heat-tree comparisons (Figure S5), we observe the largest number of heat-tree-tip differences between any two land-cover types. In other words, there are multiple species that serve as indicators of cropland (or in the case of the *Mycetophila* OTU, an indicator of Japanese cedar and eucalyptus).

## 4 DISCUSSION

To recap, our most important question is whether native forest sustains a greater biodiversity of arthropods over any of the GFGP-plantations, as we had previously found for birds (Hua et al., 2016). Our second question is whether mixed plantations can boost biodiversity over monocultures. On the second question, our previous study had found contradictory results, in that non-breeding bird diversity is higher in mixed plantations, while bee diversity is not boosted at all. To answer these two questions, we metabarcoded the arthropod bycatch of the pan traps that had originally been used to catch the bees.

### The importance of native forest

Firstly, we find that all three estimators strongly support the conclusion that native forest supports the highest levels of arthropod species richness and diversity (Figures 3, 4, 5) and that most of those species are unique to native forest (Figure 8), results that are consistent with the patterns of bird diversity that were reported in Hua et al. (2016). We thus re-affirm the biodiversity value of native forest over all GFGP-plantations and reinforce our policy recommendation of encouraging the retention and expansion of native forest in the Grain-for-Green program.

### The importance of mixed plantations

Secondly, we find that mixed plantations support levels of arthropod species richness and diversity that are higher than at least one, and possibly all three, of the monocultures (Figures 3, 4, 5). Mixed plantations are also compositionally most similar to native forests (Figure 7), and the two land-cover types uniquely share the most species (Figure 8). These results contradict the previous study’s bee-only results (Hua et al., 2016), which found no diversity boost from mixed plantations and no additional similarity of mixed plantations to native forest, but they are consistent to our previous study’s bird results, which showed higher species richness in mixed plantations over monocultures for non-breeding birds and a slightly greater degree of shared species between native forest and mixed plantations for forest-dependent birds. In short, mixed plantations not only support a higher diversity of non-breeding birds but also appear to provide a small but detectable biodiversity boost for arthropods. We thus also support the seemingly simple, but second-best, policy adjustment of encouraging checkboard plantings of different species of plantation trees as a method for boosting the biodiversity of GFGP plantations, at least in western China where we conducted this study. The value of metabarcoding is that our conclusions and policy prescriptions are now also based on a dataset that includes 536 species-resolution taxa, ranging across the Arthropoda.

Why are bees such an exception? We hypothesize here that bees are, more than the other arthropods in the pan traps, dependent on the availability of floral nectar and pollen (Roulston & Goodell, 2011). However, ferns and grasses dominate the herbaceous layer of the GFGP monocultures and mixed plantations, and flowering plants are almost absent, in part because the understory is cleared by the owning households. In contrast, as we described in *Results*, most of the 536 arthropod species in our metabarcode dataset are not directly dependent on floral nectar and pollen as dietary resources (Figure 10).

Interestingly, we found that compositional differences amongst forest types are almost entirely dominated by species turnover, not nestedness, meaning that some species were only detected in the monocultures (Figures 8, 9, S4), which provide larger stands of the same tree species relative to mixed plantations. One interpretation is that overall landscape biodiversity (gamma diversity) would be maximized by including stretches of monocultures in the landscape, but the more parsimonious explanation is that these apparently monoculture-specific species are rare, and thus less likely to be detected, in more diverse forests. For now, we cannot differentiate these two explanations.

Why should mixed plantations support more diverse arthropod and (non-breeding) bird communities? The answer is not that mixed plantations support arthropod species from cropland (Figures 7, 8). The simplest explanation is that the different tree monocultures complement each other in resource availability, probably across the seasons, such that a mosaic of monocultures - which is what a mixed plantation represents - allows more species of arthropods and birds to persist. This could be tested by sampling different parts of mixed plantations over a year, to test if the arthropod and bird communities migrate around and/or consume different plant resources. Another, non-mutually exclusive possibility is that mixed plantations might allow a more diverse vegetative understory because the monocultures differ in height and structure and thus allow sunlight to penetrate from different angles, as opposed to the uniform canopies of monocultures. A more diverse, and presumably higher-biomass, arthropod community in turn could also support a richer bird community, so our results suggest a reasonable mechanism for why bird diversity is boosted in mixed plantations.

### Metabarcoding, alpha diversity, and beta diversity

Metabarcoding provides an efficient method for interrogating diverse biodiversity samples, such as the pan-trap samples that we use here, but because of its reliance on PCR, metabarcoding datasets tend to contain a non-trivial amount of noise. This noise manifests as a large number of false-positive OTUs, which are filtered out heuristically. Such false OTUs especially complicate efforts to estimate alpha diversity. Here, we applied several filtering steps to remove false OTUs, and we also used ‘iNextPD’ to generate robust comparisons of alpha diversity by estimating phylogenetic diversity instead of species richness or diversity. (In Yu et al. (2012), we showed that metabarcoding can accurately recover the phylogenetic diversities of mock samples of known composition.)

Another approach, which became available only after we had completed the wet-lab portion of our study, is to subject each sample to multiple, independently tagged PCRs (typically three) and to bioinformatically filter out sequences that fail to appear in at least two of the PCRs above some minimum number of reads; such sequences are more likely to be PCR or sequencing errors. This is implemented in the DAMe protocol of Zepeda-Mendoza et al. (2016, also see Alberdi, Aizpurua, Gilbert, & Bohmann, 2018).

That said, we follow Magurran et al. (2015; Magurran, 2016) in recommending that biodiversity studies should focus less on explaining changes in alpha diversity and more on explaining changes in taxonomic *composition* as a function of natural and anthropogenic causes. One important reason is that anthropogenically disturbed communities can maintain species richness and diversity, even as local, or even endemic, species go extinct and are replaced by cosmopolitan species. Fortunately, metabarcoding is well suited to estimating changes in composition. For instance, Xiong and Zhan (2018) have shown that beta diversity estimates are robust to different OTU-clustering thresholds. In our study, cropland supports a species diversity similar to that mixed plantations and also just below native forest (Figures 4, 5), but cropland species are compositionally distinct from native-forest species (Figure 7), so cropland expansion cannot compensate for loss of native-forest-dependent biodiversity. Instead, our new analyses support our previous conclusions that (unsurprisingly) native forests support the most forest-dependent biodiversity and that checkboard plantings of a few tree monocultures result in the arthropod community composition of plantations becoming more similar to native forest.

## Data Accessibility

Sequence data have been submitted to GenBank under accession number SAMN09981891. Bioinformatic scripts are in Supplementary Information. R scripts and data tables are available at https://github.com/dougwyu/Sichuan2014.

## Appendix S1

**Figure S1.**
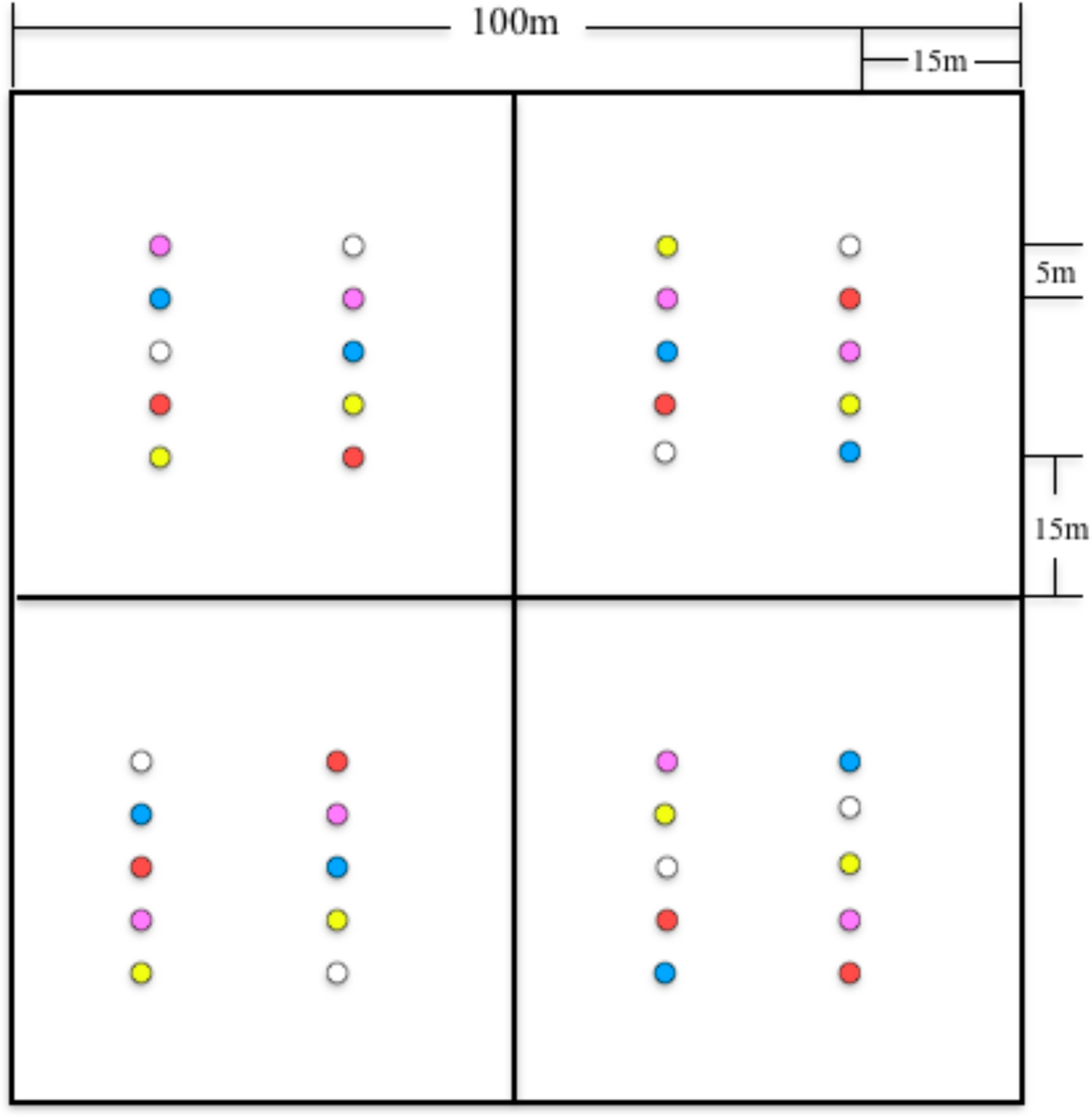
Spatial arrangement of pan traps in each one-hectare quadrat (= 1 sampling site). Each quadrat was subdivided into four subquadrats to balance pan colors. Each dot’s color represents that pan-trap’s color (white, yellow, blue, red, purple), and within each subquadrat were arranged randomly.

**Figure S2.**
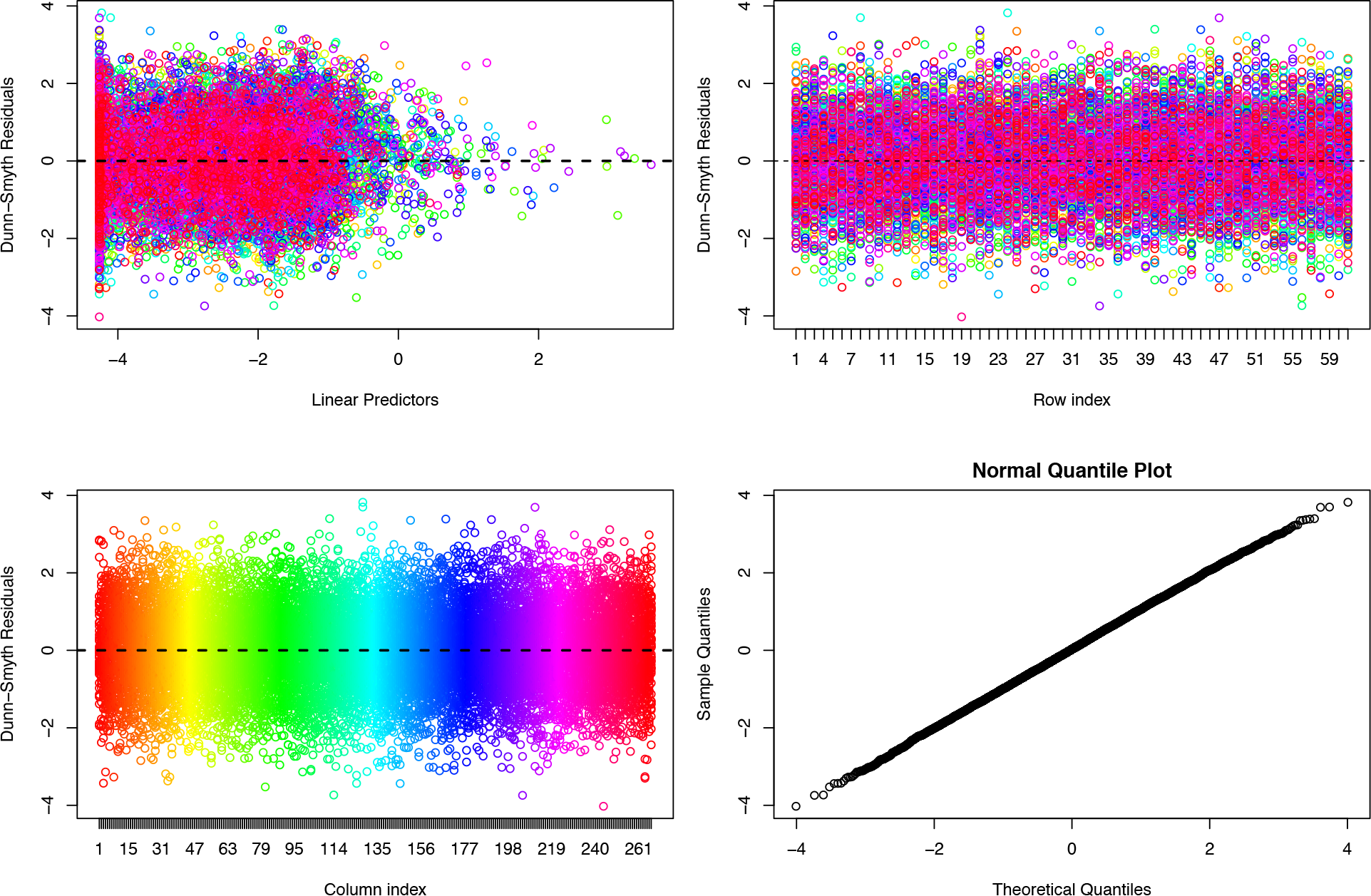
Residual plots of the *boral* model that we fit in Fig. 7.

**Figure S3.**
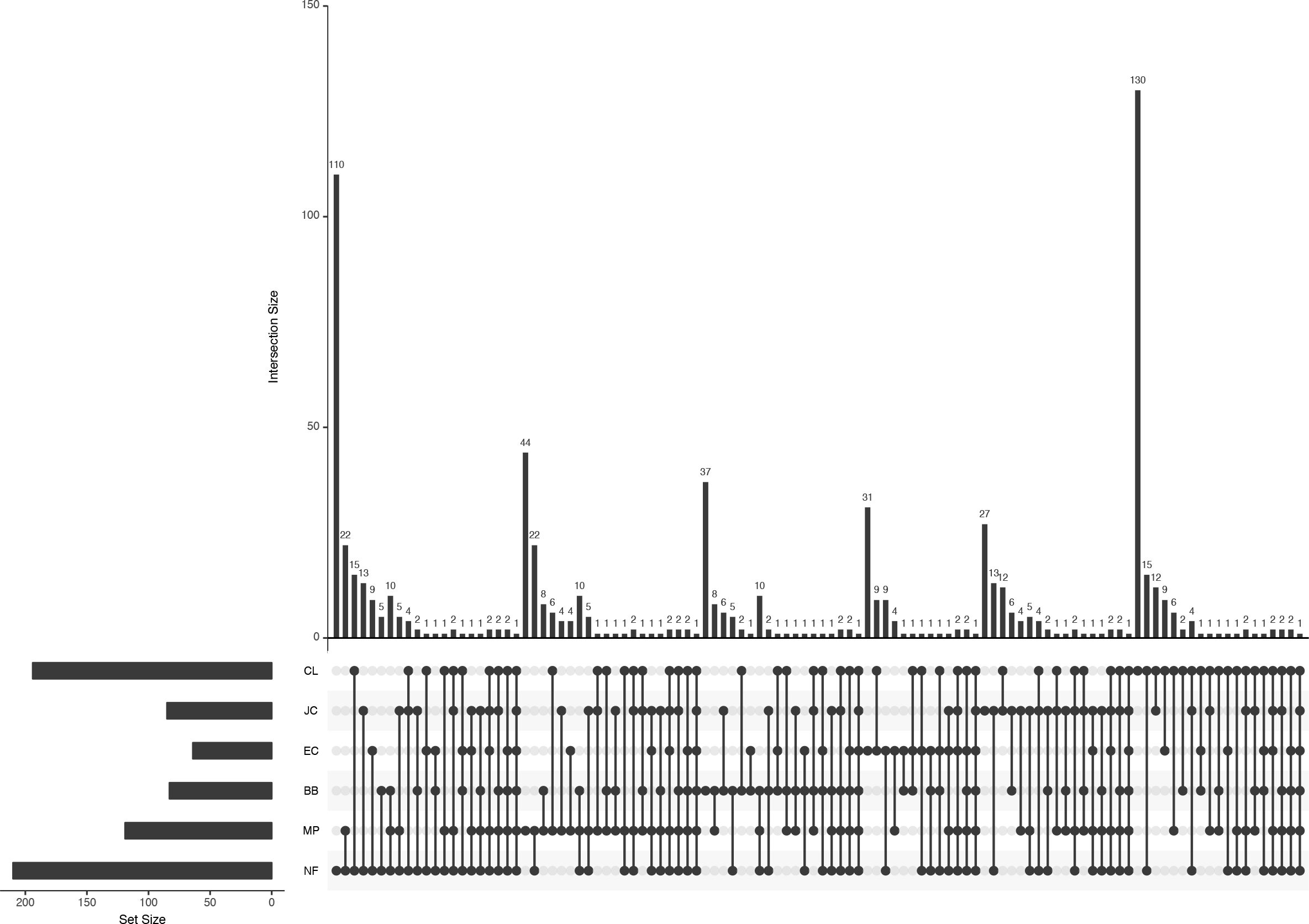
UpSetR intersection map of OTU distribution by land-cover type. Number of comparisons not truncated. Horizontal bars on the left bottom indicate the number of OTUs in each land-cover type, and vertical bars indicate the number of unique or shared OTUs. Codes for land-cover types as in Figure 1.

**Figure S4.**
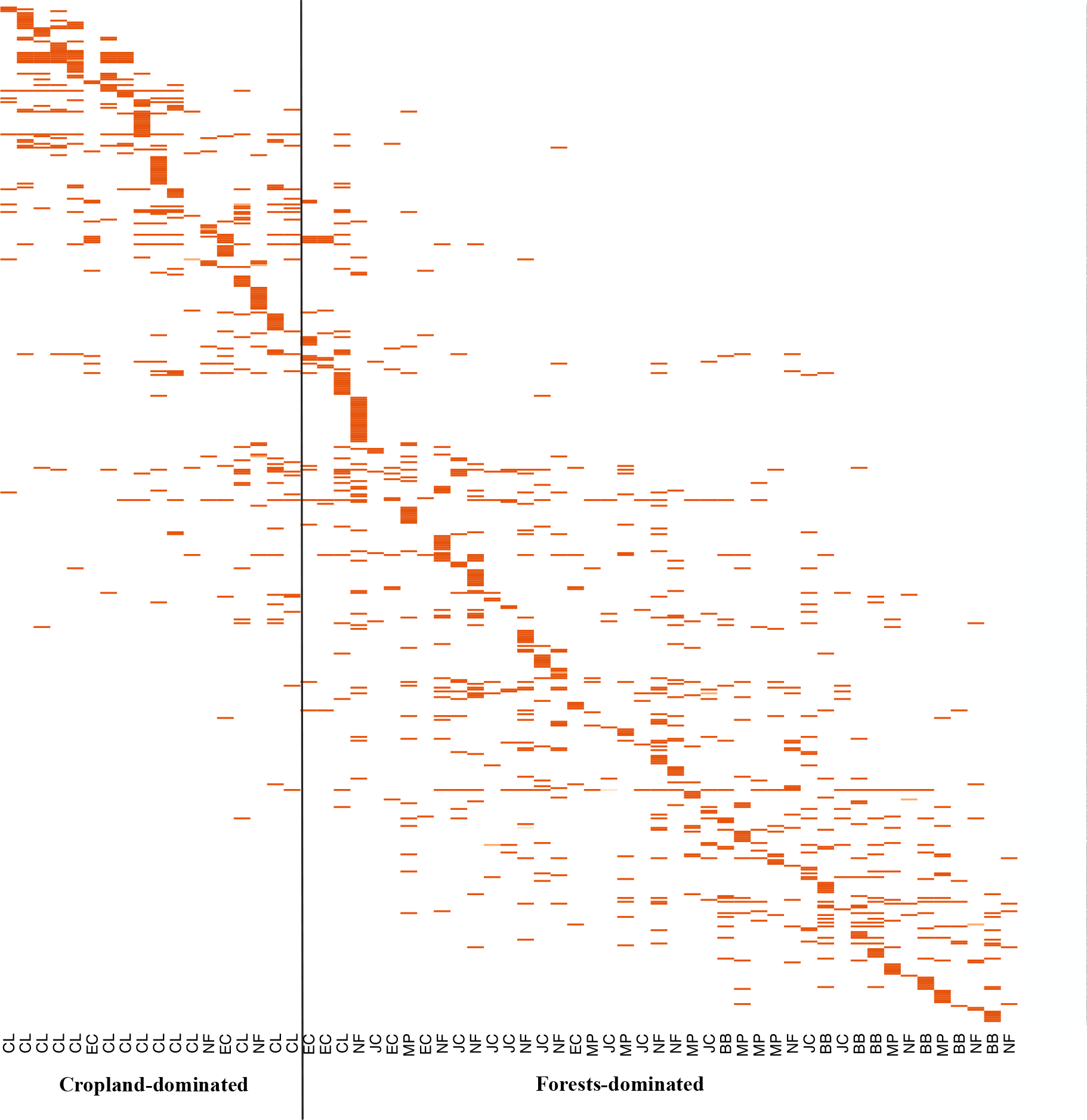
Heat map of OTU distribution by habitat type. Beta diversity is dominated by species turnover rather than by nestedness. The vertical line separates two compartments of communities, one dominated by cropland and one dominated by forest and plantations. Each column is a sample site, and rows are OTUs. Codes for land-cover types as in Figure 1.

**Figure S5.**
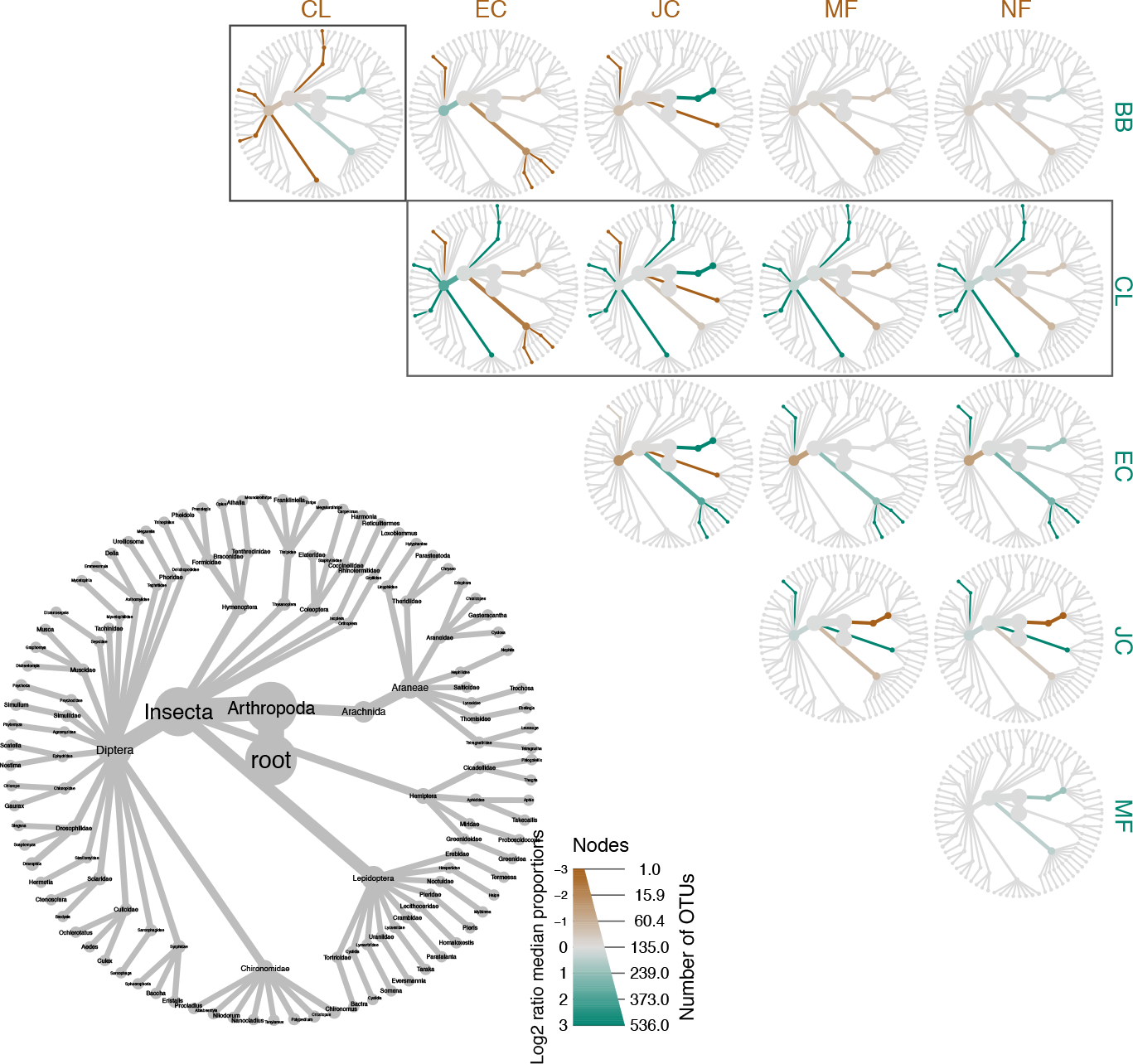
Pairwise taxonomic comparisons of all land-cover types. Interpretation the same as in Figure 10 except that cropland is included in this version of the figure (boxes). Upper right triangle: greener branches indicate taxa that are relatively more abundant (in terms of numbers of OTUs) in the land-cover type along the right column, and browner branches indicate taxa that are relatively more abundant in the land-cover type along the top row. Lower left: taxonomic identities of the branches. Note that this is a taxonomic tree, not a phylogenetic tree. Legend: width indicates number of OTUs at a given taxonomic rank, and color indicates relative differences in log_2_(number of OTUs). Codes for land-cover types as in Figure 1.

**Table S1.**
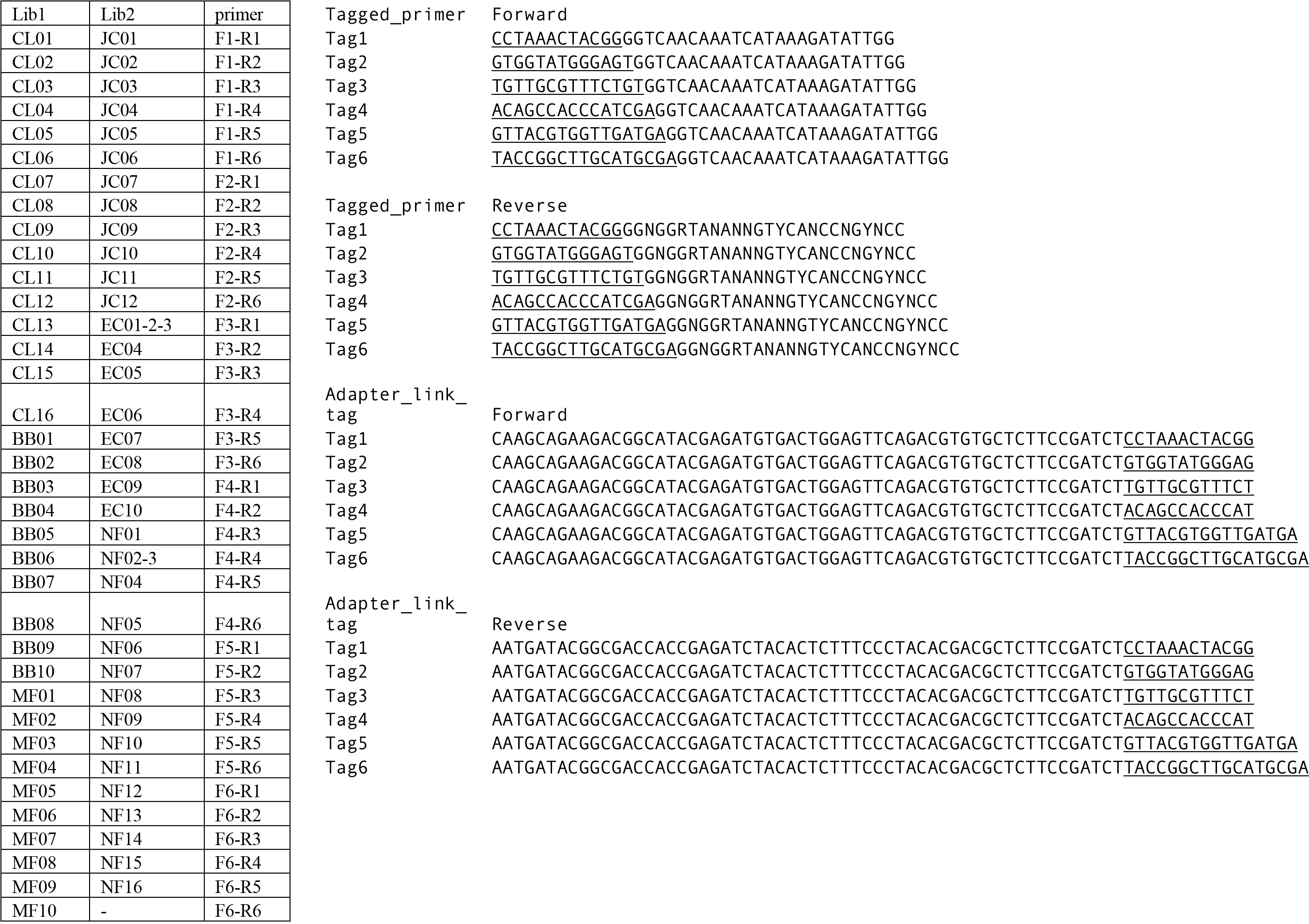
Tags and primers used in lab work and all samples relative to tagged primer. *Note*. We tailed both forward and reverse primers with sample-identifying tags, underlined above.

## REFERENCES

Alberdi, A., Aizpurua, O., Gilbert, M. T. P., & Bohmann, K. (2018). Scrutinizing key steps for reliable metabarcoding of environmental samples. Methods in Ecology and Evolution, 9(1), 134–147. https://doi.org/10.1111/2041-210X.12849

Bartholomew, Chanda S. & Prowell, D. (2005). Pan Compared to Malaise Trapping for Bees (Hymenoptera: Apoidea) in a Longleaf Pine Savanna. Journal of the Kansas Entomological Society, 78(4), 390–392.

Baselga, A., & Orme, C. D. L. (2012). Betapart: An R package for the study of beta diversity. Methods in Ecology and Evolution, 3(5), 808–812. https://doi.org/10.1111/j.2041-210X.2012.00224.x

Bryan, B. A., Gao, L., Ye, Y., Sun, X., Connor, J. D., Crossman, N. D., … Hou, X. (2018). China’s response to a national land-system sustainability emergency /704/844/685 /704/172/4081 perspective. Nature, 559(7713), 193–204. https://doi.org/10.1038/s41586-018-0280-2

Chao, A. (1987). Estimating the population size for capture-recapture data with unequal catchability. Biometrics, 43(4).

Chiu, C. H., Wang, Y. T., Walther, B. A., & Chao, A. (2014). An improved nonparametric lower bound of species richness via a modified good-turing frequency formula. Biometrics, 70(3), 671–682. https://doi.org/10.1111/biom.12200

Conway, J. R., Lex, A., & Gehlenborg, N. (2017). UpSetR: An R package for the visualization of intersecting sets and their properties. Bioinformatics, 33(18), 2938–2940. https://doi.org/10.1093/bioinformatics/btx364

Cristescu, M. E. (2014). From barcoding single individuals to metabarcoding biological communities: Towards an integrative approach to the study of global biodiversity. Trends in Ecology and Evolution, 29(10), 566–571. https://doi.org/10.1016/j.tree.2014.08.001

Deiner, K., Bik, H. M., Mächler, E., Seymour, M., Lacoursière-Roussel, A., Altermatt, F., … Bernatchez, L. (2017). Environmental DNA metabarcoding: Transforming how we survey animal and plant communities. Molecular Ecology, 26(21), 5872–5895. https://doi.org/10.1111/mec.14350

Delang, Claudio O; Yuan, Z. (n.d.). China’s Grain for Green Program. Springer US.

Deng, L., Liu, G. bin, & Shangguan, Z. ping. (2014). Land-use conversion and changing soil carbon stocks in China’s “Grain-for-Green” Program: A synthesis. Global Change Biology, 20(11), 3544–3556. https://doi.org/10.1111/gcb.12508

Deng, L., Shangguan, Z. ping, & Li, R. (2012). Effects of the grain-for-green program on soil erosion in China. International Journal of Sediment Research, 27(1), 120–127. https://doi.org/10.1016/S1001-6279(12)60021-3

Edgar, R. C. (2010). Search and clustering orders of magnitude faster than BLAST. Bioinformatics, 26(19), 2460–2461. https://doi.org/10.1093/bioinformatics/btq461

Elbrecht, V., Peinert, B., & Leese, F. (2017). Sorting things out: Assessing effects of unequal specimen biomass on DNA metabarcoding. Ecology and Evolution, 7(17), 6918–6926. https://doi.org/10.1002/ece3.3192

Foster, Z. S. L., Sharpton, T., & Grunwald, N. J. (2017). MetacodeR: An R package for manipulation and heat tree visualization of community taxonomic data from metabarcoding. BioRxiv, 071019. https://doi.org/10.1101/071019

Frøslev, T. G., Kjøller, R., Bruun, H. H., Ejrnæs, R., Brunbjerg, A. K., Pietroni, C., & Hansen, A. J. (2017). Algorithm for post-clustering curation of DNA amplicon data yields reliable biodiversity estimates. Nature Communications, 8(1). https://doi.org/10.1038/s41467-017-01312-x

Hao, X., Jiang, R., & Chen, T. (2011). Clustering 16S rRNA for OTU prediction: A method of unsupervised Bayesian clustering. Bioinformatics, 27(5), 611–618. https://doi.org/10.1093/bioinformatics/btq725

Hsieh, T. C., & Chao, A. (2017). Rarefaction and extrapolation: Making fair comparison of abundance-sensitive phylogenetic diversity among multiple assemblages. Systematic Biology, 66(1), 100–111. https://doi.org/10.1093/sysbio/syw073

Hsieh, T. C., H.Ma, K., & Chao, A. (2016). iNEXT:an Package for rarefaction and extrapolation of species diversity(Hill numbers). Ecology and Evolution, 7(12), 1451–1456. https://doi.org/10.1111/2041-210X.12613

Hua, F., Wang, L., Fisher, B., Zheng, X., Wang, X., Yu, D. W., … Wilcove, D. S. (2018). Tree plantations displacing native forests: The nature and drivers of apparent forest recovery on former croplands in Southwestern China from 2000 to 2015. Biological Conservation, 222(December 2017), 113–124. https://doi.org/10.1016/j.biocon.2018.03.034

Hua, F., Wang, X., Zheng, X., Fisher, B., Wang, L., Zhu, J., … Wilcove, D. S. (2016). Opportunities for biodiversity gains under the world’s largest reforestation programme. Nature Communications, 7, 1–11. https://doi.org/10.1038/ncomms12717

Hui, F. K. C. (2016). boral - Bayesian Ordination and Regression Analysis of Multivariate Abundance Data in r. Methods in Ecology and Evolution, 7(6), 744–750. https://doi.org/10.1111/2041-210X.12514

J Gregory Caporaso, Justin Kuczynski, Jesse Stombaugh, Kyle Bittinger, Frederic D Bushman, Elizabeth K Costello, Noah Fierer, Antonio Gonzalez Peña, Julia K Goodrich, Jeffrey I Gordon, Gavin A Huttley, Scott T Kelley, D. K. (2010). QIIME allows analysis of high-throughput community sequencing data. Nat Methods, 7(5), 335–336. https://doi.org/10.1038/nmeth.f.303.QIIME

Ji, Y., Ashton, L., Pedley, S. M., Edwards, D. P., Tang, Y., Nakamura, A., … Yu, D. W. (2013). Reliable, verifiable and efficient monitoring of biodiversity via metabarcoding. Ecology Letters, 16(10), 1245–1257. https://doi.org/10.1111/ele.12162

Kampstra, P. (2008). Beanplot: A Boxplot Alternative for Visual\nComparison of Distributions. Journal of Statistical Software, 28(code snippet 1), 1–9. https://doi.org/10.18637/jss.v028.c01

Leray, M., Yang, J. Y., Meyer, C. P., Mills, S. C., Agudelo, N., Ranwez, V., … Machida, R. J. (2013). A new versatile primer set targeting a short fragment of the mitochondrial COI region for metabarcoding metazoan diversity: Application for characterizing coral reef fish gut contents. Frontiers in Zoology, 10(1), 1–14. https://doi.org/10.1186/1742-9994-10-34

Lindenmayer, D. B., Franklin, J. F., & Fischer, J. (2006). General management principles and a checklist of strategies to guide forest biodiversity conservation. Biological Conservation, 131(3), 433–445. https://doi.org/10.1016/j.biocon.2006.02.019

Liu, J., & Diamond, J. (2005). China’s environment in a globalizing world. Nature, 435(June), 1179–1186. https://doi.org/10.1038/4351179a

Liu, J., Li, S., Ouyang, Z., Tam, C., Chen, X., Liu, J., … Chen, X. (2008). Ecological and socioeconomic effects of China’s policies for ecosystem services, 105(28), 9477–9482. https://doi.org/10.1073/pnas.0706436105

Liu, J., Ouyang, Z., Pimm, S. L., Raven, P. H., Wang, X., Miao, H., & Han, N. (2003). Protecting China’s biodiversity. Science, 300(5623), 1240–1241. https://doi.org/10.1126/science.1078868

Liu, Z., & Lan, J. (2015). The Sloping Land Conversion Program in China: Effect on the Livelihood Diversification of Rural Households. World Development, 70, 147–161. https://doi.org/10.1016/j.worlddev.2015.01.004

Long, H. L., Heilig, G. K., Wang, J., Li, X. B., Luo, M., Wu, X. Q., & Zhang, M. (2006). Land Use and Soil Erosion in the Upper Reaches of the Yangtze River : Some Socio-Economic Considerations on China’S Grain-for-Green Programme, 603(37), 589–603. https://doi.org/10.1002/ldr.736

Ma, K. (2015). Biodiversity monitoring in China: from CForBio to Sino BON. Biodiversity Science, 23(1), 1–2. https://doi.org/10.17520/biods.2015025

Ma, K., Shen, X., Grumbine, R. E., & Corlett, R. (2017). China’s biodiversity conservation research in progress. Biological Conservation, 210, 1–2. https://doi.org/10.1016/j.biocon.2017.05.029

MacGregor-Fors, I., & Payton, M. E. (2013). Contrasting Diversity Values: Statistical Inferences Based on Overlapping Confidence Intervals. PLoS ONE, 8(2), 8–11. https://doi.org/10.1371/journal.pone.0056794

Machida, R. J., Leray, M., Ho, S. L., & Knowlton, N. (2017). Data Descriptor: Metazoan mitochondrial gene sequence reference datasets for taxonomic assignment of environmental samples. Scientific Data, 4(September 2016), 1–7. https://doi.org/10.1038/sdata.2017.27

Magurran, A. E. (2016). How ecosystems change. Science, 351(6272), 448–449. https://doi.org/10.1126/science.aad6758

Magurran, A. E., Dornelas, M., Moyes, F., Gotelli, N. J., & McGill, B. (2015). Rapid biotic homogenization of marine fish assemblages. Nature Communications, 6, 1–5. https://doi.org/10.1038/ncomms9405

Mahé, F., Rognes, T., Quince, C., de Vargas, C., & Dunthorn, M. (2015). Swarm v2: highly-scalable and high-resolution amplicon clustering. PeerJ, 3, e1420. https://doi.org/10.7717/peerj.1420

Masella, A. P., Bartram, A. K., Truszkowski, J. M., Brown, D. G., & Neufeld, J. D. (2012). PANDAseq: Paired-end assembler for illumina sequences. BMC Bioinformatics, 13(1), 31. https://doi.org/10.1186/1471-2105-13-31

McMurdie, P. J., & Holmes, S. (2013). Phyloseq: An R Package for Reproducible Interactive Analysis and Graphics of Microbiome Census Data. PLoS ONE, 8(4). https://doi.org/10.1371/journal.pone.0061217

Nichols, R. V, Vollmers, C., Newsom, L. A., Wang, Y., Heintzman, P. D., Leighton, M., … Shapiro, B. (2018). Minimizing polymerase biases in metabarcoding. Molecular Ecology Resources, (February). https://doi.org/10.1111/1755-0998.12895

Nikolenko, S. I., Korobeynikov, A. I., & Alekseyev, M. A. (2013). BayesHammer: Bayesian clustering for error correction in\nsingle-cell sequencing. BMC Genomics, 14 Suppl 1(Suppl 1), S7. https://doi.org/10.1186/1471-2164-14-S1-S7

Ouyang, Z., Zheng, H., Xiao, Y., Polasky, S., Liu, J., Xu, W., … Daily, G. C. (2016). Improvements in ecosystem servcies from investments in natural capital. Science, 352(6292), 1455–1460. Retrieved from http://www.gdrc.org/sustdev/concepts/26-nat-capital.html

Piñol, J., Mir, G., Gomez-Polo, P., & Agustí, N. (2015). Universal and blocking primer mismatches limit the use of high-throughput DNA sequencing for the quantitative metabarcoding of arthropods. Molecular Ecology Resources, 15(4), 819–830. https://doi.org/10.1111/1755-0998.12355

Pyne, K. (2013). Conserving China’s Biodiversity, 3(1).

Ratnasingham, S., & Hebert, P. D. N. (2007). BARCODING, BOLD : The Barcode of Life Data System (www.barcodinglife.org). Molecular Ecology Notes, 7(April 2016), 355–364. https://doi.org/10.1111/j.1471-8286.2006.01678.x

Ren, G., Young, S. S., Wang, L., Wang, W., Long, Y., Wu, R., … Yu, D. W. (2015). Effectiveness of China’s National Forest Protection Program and nature reserves. Conservation Biology, 29(5), 1368–1377. https://doi.org/10.1111/cobi.12561

Rognes, T., Flouri, T., Nichols, B., Quince, C., & Mahé, F. (2016). VSEARCH: a versatile open source tool for metagenomics. PeerJ, 4, e2584. https://doi.org/10.7717/peerj.2584

Roulston, T. H., & Goodell, K. (2011). The Role of Resources and Risks in Regulating Wild Bee Populations. Annual Review of Entomology, 56(1), 293–312. https://doi.org/10.1146/annurev-ento-120709-144802

Sayer, J. A., & Sun, C. (2003). Impacts of policy reforms on forest environments and biodiversity. China’s Forests: Global Lessons from Market Reforms, 177–194. https://doi.org/10.4324/9781936331239

Schirmer, M., Ijaz, U. Z., D’Amore, R., Hall, N., Sloan, W. T., & Quince, C. (2015). Insight into biases and sequencing errors for amplicon sequencing with the Illumina MiSeq platform. Nucleic Acids Research, 43(6). https://doi.org/10.1093/nar/gku1341

Schnell, I. B., Bohmann, K., & Gilbert, M. T. P. (2015). Tag jumps illuminated - reducing sequence-to-sample misidentifications in metabarcoding studies. Molecular Ecology Resources, 15(6), 1289–1303. https://doi.org/10.1111/1755-0998.12402

Schubert, M., Lindgreen, S., & Orlando, L. (2016). AdapterRemoval v2: Rapid adapter trimming, identification, and read merging. BMC Research Notes, 9(1), 1–7. https://doi.org/10.1186/s13104-016-1900-2

Shen, Y., Liao, X., & Yin, R. (2006). Measuring the socioeconomic impacts of China’s Natural Forest Protection Program. Environment and Development Economics, 11(6), 769–788. https://doi.org/10.1017/S1355770X06003263

Stamatakis, A. (2014). RAxML version 8: A tool for phylogenetic analysis and post-analysis of large phylogenies. Bioinformatics, 30(9), 1312–1313. https://doi.org/10.1093/bioinformatics/btu033

Tao, Y., Huang, D., Jin, X., & Guo, L. (2010). Make an Inventory of China’s Biodiversity. Bulletin of the Chinese Academy of Sciences, 24(4), 212–217.

Wang, J., Peng, J., Zhao, M., Liu, Y., & Chen, Y. (2017). Significant trade-off for the impact of Grain-for-Green Programme on ecosystem services in North-western Yunnan, China. Science of the Total Environment, 574, 57–64. https://doi.org/10.1016/j.scitotenv.2016.09.026

Wang, Q., Garrity, G. M., Tiedje, J. M., & Cole, J. R. (2007). Naïve Bayesian classifier for rapid assignment of rRNA sequences into the new bacterial taxonomy. Applied and Environmental Microbiology, 73(16), 5261–5267. https://doi.org/10.1128/AEM.00062-07

Wang, Y., Naumann, U., Wright, S. T., & Warton, D. I. (2012). Mvabund- an R package for model-based analysis of multivariate abundance data. Methods in Ecology and Evolution, 3(3), 471–474. https://doi.org/10.1111/j.2041-210X.2012.00190.x

Wang, Z. J., Jiao, J. Y., Rayburg, S., Wang, Q. L., & Su, Y. (2016). Soil erosion resistance of “Grain for Green” vegetation types under extreme rainfall conditions on the Loess Plateau, China. Catena, 141, 109–116. https://doi.org/10.1016/j.catena.2016.02.025

Warton, D. I., Blanchet, F. G., O’Hara, R. B., Ovaskainen, O., Taskinen, S., Walker, S. C., & Hui, F. K. C. (2015). So Many Variables: Joint Modeling in Community Ecology. Trends in Ecology and Evolution, 30(12), 766–779. https://doi.org/10.1016/j.tree.2015.09.007

Wei, F., Swaisgood, R., Hu, Y., Nie, Y., Yan, L., Zhang, Z., … Zhu, L. (2015). Progress in the ecology and conservation of giant pandas. Conservation Biology, 29(6), 1497–1507. https://doi.org/10.1111/cobi.12582

Wei, Y., Yu, D., Lewis, B. J., Zhou, L., Zhou, W., Fang, X., … Dai, L. (2014). Forest carbon storage and tree carbon pool dynamics under natural forest protection program in northeastern China. Chinese Geographical Science, 24(4), 397–405. https://doi.org/10.1007/s11769-014-0703-4

Xie Gaodi, Cao Shuyan, Yang Qisen, Xia Lin, Fan Zhiyong, Chen Boping, Zhou Shuang, Chang Youde, Ge Liqiang, Seth Cook, S. H. (2012). China Ecological Footprint Report 2012, 64.

Xiong, W., & Zhan, A. (2018). Testing clustering strategies for metabarcoding-based investigation of community-environment interactions. Molecular Ecology Resources, (July), 1–13. https://doi.org/10.1111/1755-0998.12922

Xu, Haigen; Wang, Shunqing; Xue, D. (1999). Biodiversity conservation in China:\rlegislation, plans and measures. Biodiversity and Conservation 8: 819–837, 1999, 819–837.

Xu, J., Yin, R., Li, Z., & Liu, C. (2006). China’s ecological rehabilitation: Unprecedented efforts, dramatic impacts, and requisite policies. Ecological Economics, 57(4), 595–607. https://doi.org/10.1016/j.ecolecon.2005.05.008

Yin, R., Liu, C., Zhao, M., Yao, S., & Liu, H. (2014). The implementation and impacts of China’s largest payment for ecosystem services program as revealed by longitudinal household data. Land Use Policy, 40, 45–55. https://doi.org/10.1016/j.landusepol.2014.03.002

Yin, R., Yin, G., & Li, L. (2009). Assessing China’s ecological restoration programs: What’s been done and what remains to be done? An Integrated Assessment of China’s Ecological Restoration Programs, 21–38. https://doi.org/10.1007/978-90-481-2655-2_2

Yu, D. W., Ji, Y., Emerson, B. C., Wang, X., Ye, C., Yang, C., & Ding, Z. (2012). Biodiversity soup: Metabarcoding of arthropods for rapid biodiversity assessment and biomonitoring. Methods in Ecology and Evolution, 3(4), 613–623. https://doi.org/10.1111/j.2041-210X.2012.00198.x

Zepeda-Mendoza, M. L., Bohmann, K., Carmona Baez, A., & Gilbert, M. T. P. (2016). DAMe: A toolkit for the initial processing of datasets with PCR replicates of double-tagged amplicons for DNA metabarcoding analyses. BMC Research Notes, 9(1), 1–13. https://doi.org/10.1186/s13104-016-2064-9

Zhai, D. L., Xu, J. C., Dai, Z. C., Cannon, C. H., & Grumbine, R. E. (2014). Increasing tree cover while losing diverse natural forests in tropical Hainan, China. Regional Environmental Change, 14(2), 611–621. https://doi.org/10.1007/s10113-013-0512-9

Zheng, H., & Cao, S. (2014). Threats to China???s Biodiversity by Contradictions Policy. Ambio, 44(1), 23–33. https://doi.org/10.1007/s13280-014-0526-7

Zhou, H., Van Rompaey, A., & Wang, J. (2009). Detecting the impact of the “Grain for Green” program on the mean annual vegetation cover in the Shaanxi province, China using SPOT-VGT NDVI data. Land Use Policy, 26(4), 954–960. https://doi.org/10.1016/j.landusepol.2008.11.006

